# *Staphylococcus aureus* GpsB, Stk, and VraSR form a cell wall surveillance network to maintain cell envelope integrity and mediate antibiotic resistance

**DOI:** 10.64898/2026.10.08.757784

**Authors:** Lily J. McKnight, Olivia Payne, Asher King, Jason Carroll, Briana Boyer, Andrew Brown, Lauren R. Hammond, Prahathees J. Eswara

**Author notes:** Address correspondence to Prahathees J. Eswara.

## Abstract

Antibiotic resistance is an alarming threat to public health worldwide. Specifically, drug-resistant *Staphylococcus aureus* remains a major contributor to global mortality. Therefore, in-depth understanding of antibiotic resistance mechanisms is urgently needed to inform alternative therapeutic strategies. We previously showed that GpsB is a critical regulator of cell wall synthesis in *S. aureus*. GpsB executes its regulatory role via protein-protein interactions. Yet, how GpsB function is modulated in this organism has remained unclear. Intriguingly, a prior proteomics report indicated that GpsB is phosphorylated at six different residues by the sole serine/threonine kinase Stk. Hence, we explored the physiological significance of GpsB phosphorylation. We identify three phosphosites that are each independently sufficient to drive GpsB phosphoregulation, which likely alters its partner preference. We find that cells lacking *gpsB* exhibit heightened fosfomycin sensitivity, and that this is exacerbated by *stk* deletion. Our findings reveal that GpsB collaborates with Stk and the VraSR cell wall stress response system to coordinate the demand-driven availability of cell wall precursors. Collectively, this report establishes GpsB as a pivotal peptidoglycan supply chain sentinel that together with Stk and VraSR forms an integrated cell wall stress monitoring network that maintains cell envelope homeostasis and aids in antibiotic resistance.

**Significance:** The combination of multidrug resistance and an extensive arsenal of virulence factors makes *Staphylococcus aureus* a formidable pathogen. Therefore, to help devise novel therapeutic strategies a deeper understanding of the molecular mechanisms underlying antibiotic resistance is critical. In this report, we reveal that GpsB plays an important role in cell wall stress response. We find that the Stk-dependent serine/threonine phosphorylation of GpsB influences its interaction partner preference. Additionally, our results show that loss of *gpsB* sensitizes cells to fosfomycin-induced cell wall stress, and that this effect is further exacerbated by *stk* deletion. Furthermore, we uncover previously unrecognized GpsB interaction partners involved in cell wall precursor biosynthesis. These findings elucidate the molecular basis underlying the role of GpsB in ensuring cell envelope integrity. Taken together, our study establishes GpsB as a critical mediator of cell wall stress response and antibiotic resistance.

## Introduction

*Staphylococcus aureus* is a human-adapted pathogen and the leading cause of bacterial infection associated deaths worldwide due to its virulence potential and antibiotic resistance traits (1, 2). Commercial antibiotics that are commonly used to treat *S. aureus* infections target the cell envelope (3, 4). Our previous studies elucidated the role of *S. aureus* GpsB in cell division and cell wall synthesis (5–7). The crucial role of GpsB in regulating cell division, peptidoglycan synthesis, and other cell envelope related (teichoic acids) pathways in *S. aureus* and other closely related organisms has been well established (8–12). Remarkably, GpsB influences these diverse processes through directly interacting with multiple proteins. Notable partners of GpsB include the central cell division protein FtsZ (5, 7, 13); penicillin binding proteins PBP2 and PBP4 (7, 14, 15); and wall teichoic acids related proteins TarO and TarG (6, 14). Additionally, GpsB homologs in other organisms are subject to serine/threonine kinase mediated phosphoregulation (16–21). A proteome-wide phosphorylation analysis in *S. aureus* identified six different S/T phosphosites in GpsB (22). In *S. aureus*, Stk (PknB1 or Stk1) is the only S/T kinase in most strain types and has been linked to cell wall synthesis, metabolism, and antibiotic resistance (23–27). Yet, how *S. aureus* GpsB is influenced by phosphorylation and whether it plays any role in antibiotic resistance has remained unexplored. Therefore, we aimed to address this knowledge gap.

In this article, using systematic site-directed mutagenesis to mimic either phosphorylated (S/T E) or un/de-phosphorylated (S/T A) state, we show that GpsB phosphoregulation is primarily mediated by three phosphosites independently. With the aid of the bacterial two-hybrid (BTH) assay, we reveal that phosphomutation does not affect self-interaction. Yet, remarkably, it does influence its partner choice. Briefly, we show that phosphomimetic GpsB shows altered high and low preference for Stk and PBP2, respectively. Intriguingly, this preference is reversed with phosphoablative mutation. Next, we tested the sensitivity of cells lacking *gpsB*, *stk*, or both to the cell wall targeting antibiotic fosfomycin (Fos). Our results uncovered that the Δ*stk* strain is more sensitive to Fos than wild type (WT) control. In contrast, we see that the *gpsB* deletion strain is even more sensitive than the *stk* null mutant. This data shows that GpsB plays a bigger role than Stk in mediating Fos resistance. Additionally, we see that cells lacking both *stk* and *gpsB* are extremely sensitive to Fos. This, together with phosphomutant complementation experiments, reveal that GpsB phosphoregulation is critical for optimal response to Fos.

The main target of Fos is MurA, the primary housekeeping enzyme responsible for cell wall precursor synthesis (28, 29). Upon experiencing cell wall stress, *S. aureus* selectively increases the expression of *murZ* (29). Consequently, MurA together with its homolog MurZ elevate the peptidoglycan (PG) precursor level in response to cell wall stress (29). It is to be noted that both MurA and MurZ are inhibited by Fos (29). Multiple lines of evidence support the role of S/T kinase in influencing MurA stability in *S. aureus* and other related organisms. Briefly, the protease adapters ReoM and ReoY work together to turnover MurA, and Stk-dependent ReoM phosphorylation inhibits this process and leads to MurA accumulation (18, 30, 31). Curiously, in other species, ReoM (IreB) is known to directly interact with GpsB (21) and/or S/T kinase-mediated phosphorylation of ReoM was linked to GpsB (20, 32–34). Separately, Stk through phosphorylation also hinders the DNA binding ability of the two-component system (TCS) response regulator (VraR) (35), which is responsible for *murZ* upregulation in *S. aureus* (36). Thus, the Stk pathway seems to prioritize MurA over MurZ. As our data indicates that absence of GpsB, but not Stk, leads to increased Fos sensitivity, we hypothesized that GpsB must mediate Fos resistance directly through MurA and/or MurZ independent of Stk. Our BTH assay shows that GpsB directly interacts with MurA and MurZ, as well as ReoM and ReoY. Moreover, we demonstrate that Δ*gpsB* Fos hypersensitivity is reversed by additional deletion of *reoY*, which is expected to allow MurA accumulation. Using a luciferase-based *murZ* transcriptional reporter, we show that unlike cells lacking *stk*, the *gpsB* null mutant is defective in upregulating *murZ* in the presence of Fos. Therefore, we probed the link between GpsB and the VraSR TCS through BTH assay. Our results reveal that GpsB is indeed able to directly interact with both VraS and VraR.

In summary, this report elucidates that GpsB: (i) is critical to alleviate cell wall stress; (ii) phosphoregulation by Stk alters its partner preference; (iii) reinforces PG precursor production by directly interacting with VraSR TCS components; and (iv) likely also restricts MurA/MurZ proteolysis/enzymatic function through its interaction with ReoM. Overall, our model illustrates that GpsB/Stk/VraSR coordination finetunes the need-based cell wall precursor level and this ultimately is of paramount importance for survival in the presence of cell wall stress.

## Results

### The residues T21, T75, and S76 are highly critical for GpsB phosphoregulation

GpsB is a coiled-coil protein containing structured N-terminal domain (NTD) and C-terminal domain (CTD) that are connected by a highly flexible intrinsically disordered linker region (IDR) (**Fig. 1A**) (9). The NTD of GpsB forms dimer and harbors residues needed for peripheral interaction with the membrane while its CTD exists as a hexamer, more precisely as a dimer of trimers (9, 37). Uniquely, *S. aureus* GpsB (*Sa* GpsB) also has a hinge-like region within the longer alpha-helix in its NTD (**Fig. 1A**; depicted as) (7). However, owing to its IDR, the high-resolution structure of full-length GpsB in its hexameric state remains unsolved. It is also established that GpsB via its NTD (**Fig. 1**; yellow rectangles) recognizes many of its substrates containing a signature motif (R-X-X-R) with additional stringency conferred by adjoining residues (7, 9, 12, 13, 38).

**Fig. 1:**
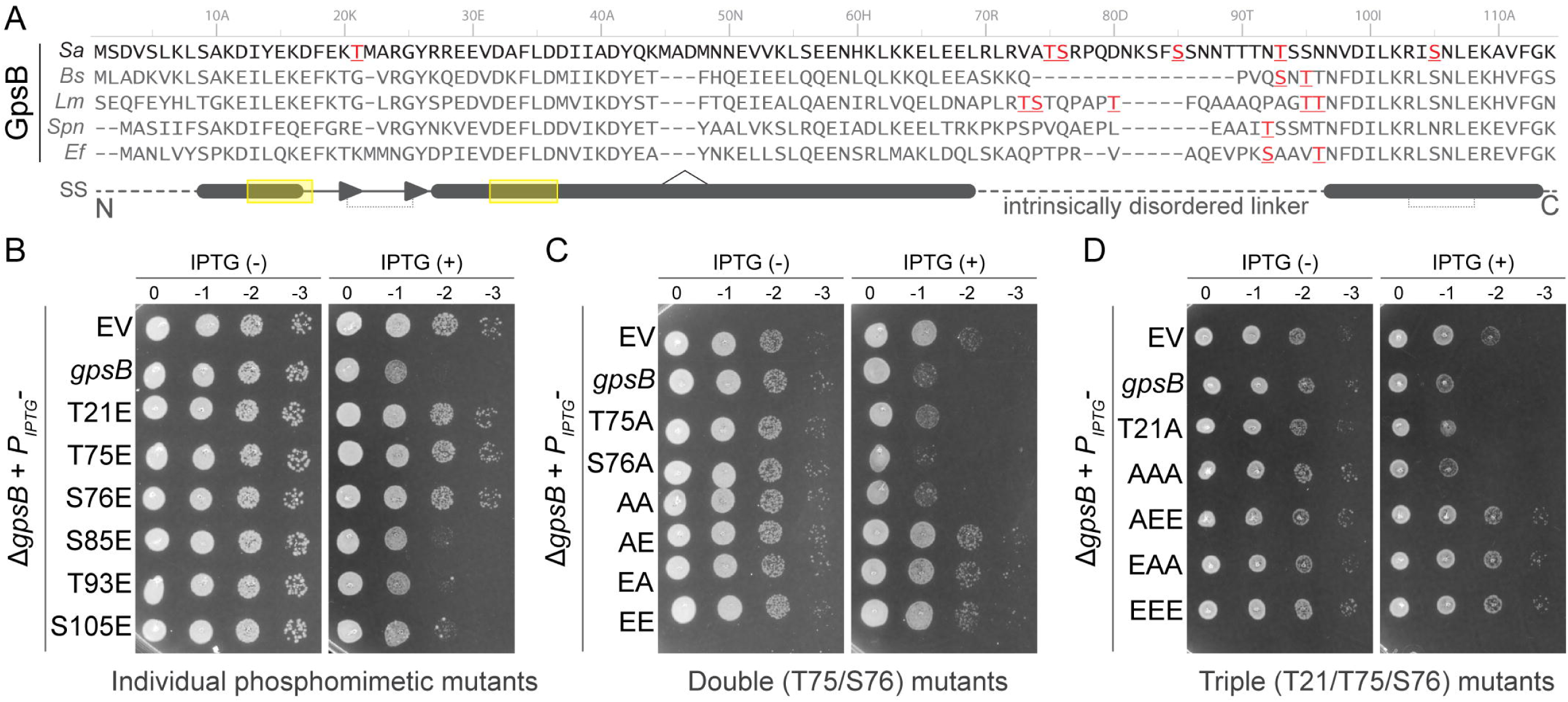
Three unique S/T residues are individually sufficient for GpsB phosphoregulation. **(A)** Multiple sequence alignment of GpsB from *S. aureus* (*Sa*), *B. subtilis* (*Bs*), *L. monocytogenes* (*Lm*), *S. pneumoniae* (*Sp*), *and E. faecalis* (*Ef*). Known S/T phosphosites are shown in red font. The secondary structure (SS) features including alpha helices (filled cylinders), β-hairpin-like loop (filled triangles), and disordered regions (dashed lines) are indicated. The residues that mediate inter-subunit interactions (dotted lines) and the partner binding interface (yellow rectangles) are also shown. The hinge-like region specific to *S. aureus* is depicted using symbol. **(B-D)** Spot titer assay showing the growth ten-fold serially diluted cultures of *S. aureus* (RN4220) Δ*gpsB* strain hosting the empty vector (EV) or the vector harboring an IPTG-inducible copy of *gpsB* or *gpsB* phosphomutants. The TSA plates without (left; -) or with (right; +) IPTG were imaged after overnight incubation at 37 °C. Strains used: EV (LM74); *gpsB* (LM75); T21E (LM235); T75E (LM236); S76E (LM237); S85E (LM238); T93E (LM239); S105E (LM240); T21A (LM241); T75A (LM242); S76A (LM243); EEE (LM244); EAA (LM245); AEE (LM246); AAA (LM247); AE (LM248); EA (LM249); EE (LM250); AA (LM251). Representative images of three independent biological replicates are shown.

Pertinent to this study, GpsB is among the several substrates of Stk in *S. aureus* (22, 27). Specifically, GpsB is phosphorylated at six different S/T residues: T21, T75, S76, S85, T93 and S105 (22). Intriguingly, GpsB phosphorylation sites in *S. aureus* and other organisms appear to be clustered within IDR (**Fig. 1A**; residues shown in red font). Additionally, *Sa* GpsB also contains phosphosites within its NTD (T21) and CTD (S105). Notably, T21 is part of the region that participates in inter-subunit interaction within the dimer, and S105 is within the R/E residues that are critical for CTD trimerization (**Fig. 1A**; depicted as dotted lines) (9).

In this study, we sought to investigate which of these 6 phosphorylatable sites are important for *Sa* GpsB post-translational regulation. For this purpose, we generated phosphomimetic (S/T E) substitutions of all 6 residues individually. Of note, these GpsB phosphomimetic mutants cannot be dephosphorylated by Stp phosphatase and permit us to examine the effects of GpsB phosphorylation albeit indirectly. Next, to probe the functionality of GpsB phosphomutants we capitalized on our prior knowledge of growth inhibition seen with *Sa* GpsB overproduction (7). Briefly in a *gpsB* null background, overexpression of *gpsB* is toxic to growth (**Fig. 1B**; second row). This is not the case in this strain when the inducer (IPTG) is absent or in wild type (WT) background. We then investigated the effects of overexpression of various phosphomimetic mutants individually or in combination. Remarkably, as shown in **Fig. 1B**, glutamate substitution of residues T21, T75, and S76 rendered them non-toxic while the remaining sites (S85, T93, and S105) appeared to be largely unaffected. Importantly, we ensured that all phosphomutants are stably produced (**Fig. S1**). Based on our observation in this experimental condition, T21, T75, and S76 appear to be the key sites of GpsB phosphomodulation.

Next, we wondered whether any one of these sites are individually sufficient for rendering phosphomimetic GpsB non-toxic upon overproduction. To dissect this, we generated phosphoablative (S/T A) mutants to prevent Stk phosphorylation of that particular site. We note that overproduction of GpsB harboring T75A or S76A mutations individually, or together (AA), does not alter its growth-inhibitory function. In striking contrast, glutamate substitution of one or both of these two sites abolishes the overproduction toxicity, as seen for T75A S76E (AE), T75E S76A (EA), or T75E S76E (EE) double mutants (**Fig. 1C**). This result revealed that phosphorylation of either T75 or S76 is likely sufficient to post-translationally control GpsB function. Finally, we also investigated the specific role of T21 in the context of other phosphosites. For this, we generated triple mutants of T21 with T75 and S76 AA/EE double mutants. As shown in **Fig. 1D**, both T21A single and T21A T75A S76A (AAA) triple mutants retain their growth inhibitory functions. Conversely, T21A T75E S76E (AEE), T21E T75E S76E (EEE), and more importantly T21E T75A S76A (EAA; compared to AA in **Fig. 1C**) are all not toxic upon overproduction. Overall, this dataset demonstrates that Stk-mediated regulation of GpsB can be solely and independently achieved by T21, T75, or S76 phosphorylation. As GpsB phosphorylation in IDR linker is the most conserved regulatory mechanism, we chose to conduct further experiments with the T75 phosphomutant.

### GpsB phosphorylation may alter its partner preference

It is known that protein phosphorylation may promote partner-switching behavior either by positively or negatively influencing the affinity of a given protein to its interaction partners (39–41). Similarly, ReoM-mediated proteolysis of MurA is abolished by Stk-mediated phosphorylation of ReoM (18, 30, 42). Therefore, we investigated whether *Sa* GpsB partner preference is altered by phosphorylation. For this, we used bacterial two-hybrid (BTH) assay (43). Briefly, positive interaction between two proteins in BTH system can be inferred by color change in the presence of a chromogenic substrate (**Fig. 2A**; see controls). Our results show that self-interaction of GpsB and phosphomutants (T75E and T75A) are unaffected (**Fig. 2A**). Next, we probed the interaction between GpsB/phosphomutants and its known partners: Stk, Stp, TarG (6), and proteins related to cell wall synthesis including PBP2 (15). In support of our prediction, we observe differential interaction with Stk and PBP2 (**Fig. 2B**, see arrowheads). Specifically, we find Stk interaction to be the strongest with T75E. On the other hand, PBP2 interaction is strongest with unmutated GpsB and this is substantially weakened when the T75 residue of GpsB is mutated irrespective of whether the mutation is phosphomimetic or phosphoablative. We infer this to mean that the availability of this threonine is critical for PBP2 recognition especially in the non-native *Escherichia coli*-based BTH system. However, this is not the case for Stp and TarG. Likewise, PBP2a, a key determinant of methicillin resistance (44), also interacted with all versions of GpsB. Notably, this is the first report to reveal the interaction between PBP2a and GpsB. On the contrary, PBP1 and PBP3 do not appear to directly interact with GpsB in this assay. Although we previously showed that PBP4 and FtsZ, a tubulin-like GTPase, are also interaction partners of GpsB using surface plasmon resonance and other biochemical and structural biology techniques (7), we were unable to reproduce this interaction in BTH assay. This might be because, unlike other partners, these two proteins interact with GpsB through their C-termini (7, 13). Thus, we conducted biochemical experiments with purified FtsZ and GpsB, as well as the function-impaired T75E mutant (**Fig. 1B**), as described previously (13). In comparison to GpsB control, we see that both the binding affinity (**Fig. 2C**) and stimulation of GTPase activity (**Fig. 2D**) are negatively affected in the T75E mutant. As T75 phosphosite in the IDR is not proximal to the partner-binding interface located in the N-terminal domain (**Fig. 1A**), our result implies that phosphorylation at this site likely induces changes in the binding pocket conformation. However, this does not affect GpsB self-interaction (**Fig. 2A**). Whether higher order oligomerization (hexamerization) of GpsB is affected by phosphorylation remains to be elucidated. Nonetheless, our data supports the notion that GpsB partner preference is altered by Stk-mediated phosphorylation at least for a subset of interaction partners. Simply, as illustrated in **Fig. 2E**, Stk-dependent GpsB phosphorylation may serve to modulate GpsB-PBP2 interaction and/or perhaps weaker GpsB-PBP2 interaction may make GpsB more available for interaction with, and stimulation of, Stk kinase. This underscores how GpsB phosphoregulation can serve as a critical toggle switch through which Stk/Stp may be able to fine-tune cell wall synthesis especially when experiencing cell envelope stress.

**Fig. 2:**
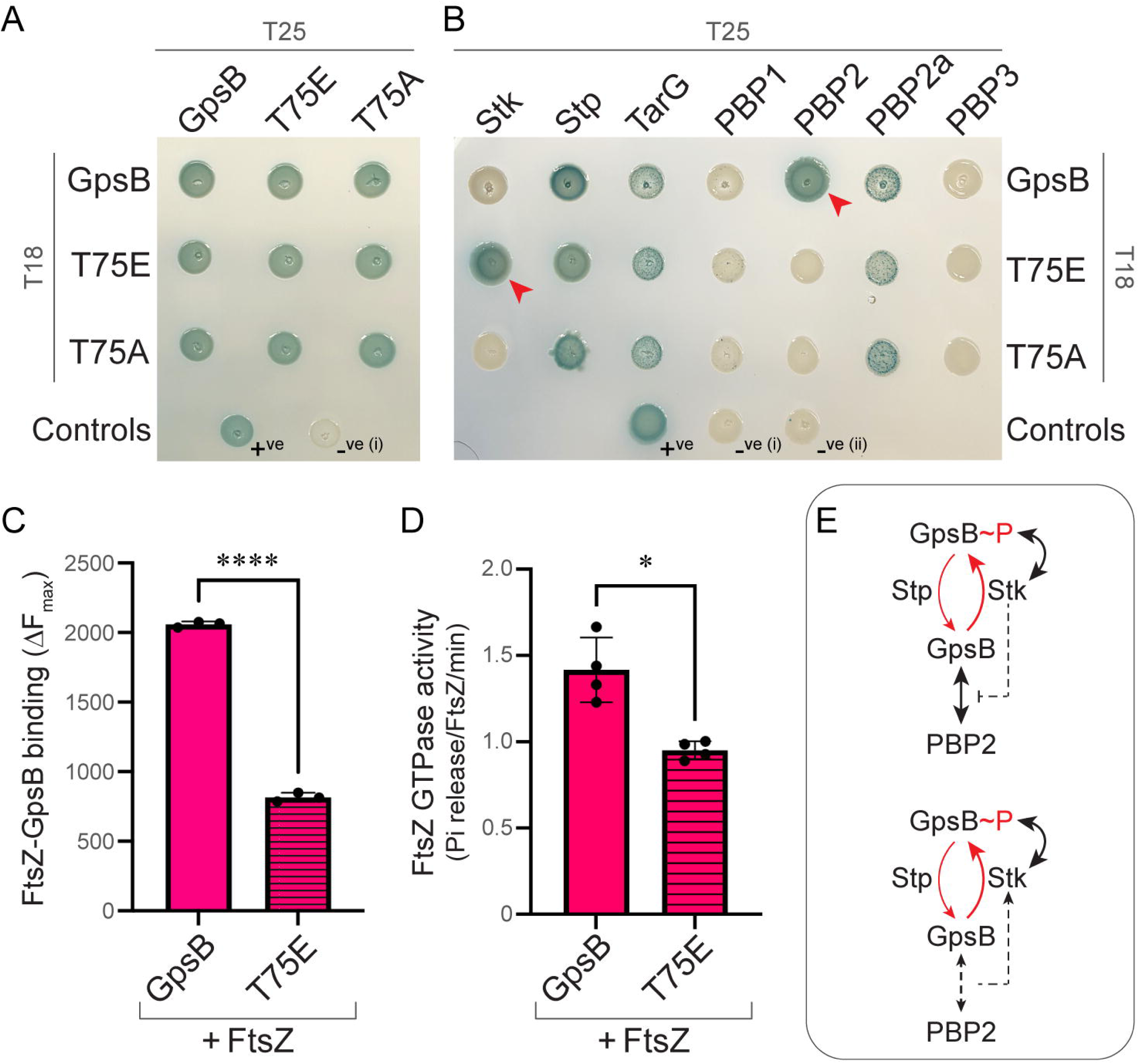
Phosphomutation influences GpsB partner preference. (**A-B**) Bacterial two-hybrid (BTH) assay results. Pair-wise self-interaction between GpsB, T75E, and T75A (panel A) and interactions between these GpsB/phosphomutants and known/putative partners (panel B) were tested in the *Escherichia coli* BTH101 strain. Briefly, positive interaction in this adenylate cyclase-dependent BTH assay produces blue color on LB agar containing the chromogenic substrate X-gal. Specific details regarding the positive and negative controls are described under methods. Representative plate images of three independent biological replicates are shown. (**C-D**) FtsZ interaction with GpsB and T75E phosphomimetic mutant were investigated using biochemical techniques. Fluorescence change in FITC-labeled FtsZ was monitored upon titration with increasing concentration of GpsB/T75E (panel C). Briefly, an increased value of change in maximal fluorescence (ΔF_max_) suggests stronger binding affinity. The GTPase activity of FtsZ was assessed in the presence of GpsB or T75E mutant (panel D). Error bars represent standard deviation, n=4. **** and * indicate *p*<0.0001 and 0.01 respectively. (**E**) Inference models showing how Stk-mediated GpsB phosphorylation may influence, or is influenced by, GpsB-PBP2 interaction.

### GpsB is a key mediator of fosfomycin resistance

Our data thus far, with the aid of phosphomutants, uncovered the potential of GpsB phosphorylation in regard to its function (**Fig. 1**) and partner choice (**Fig. 2**). However, this did not provide us with the context as to when GpsB phosphoregulation may become important. It is known that S/T kinases often participate in stress response and play a key role in maintaining cell envelope homeostasis (26, 27, 45). Thus, we investigated the relevance of GpsB and its phosphorylation in the presence of the cell wall targeting antibiotic fosfomycin (Fos). Briefly, Fos specifically inhibits MurA/MurZ, depletes the intracellular PG precursor level, and induces cell wall stress (28, 29). As shown in **Fig. 3A**, growth curves (OD_600_) of cells lacking *gpsB*, *stk*, or both grew similar to WT control in the absence of antibiotic stress. However, surprisingly, we find that cells lacking *gpsB* are more sensitive to Fos than *stk* mutant in comparison with the WT control. Furthermore, our results show that absence of both *gpsB* and *stk* is highly deleterious, especially when the cells are grown in the presence of elevated Fos concentration (5 µg/ml). This data highlights that GpsB is more important than Stk for alleviating cell wall stress caused by PG precursor depletion. However, absence of both GpsB and Stk make cells hypersensitive to Fos. Thus, in the presence of antibiotic stress and when GpsB is absent, the presence of Stk and presumably phosphorylation of its substrates become crucial for maintaining viability.

**Fig. 3:**
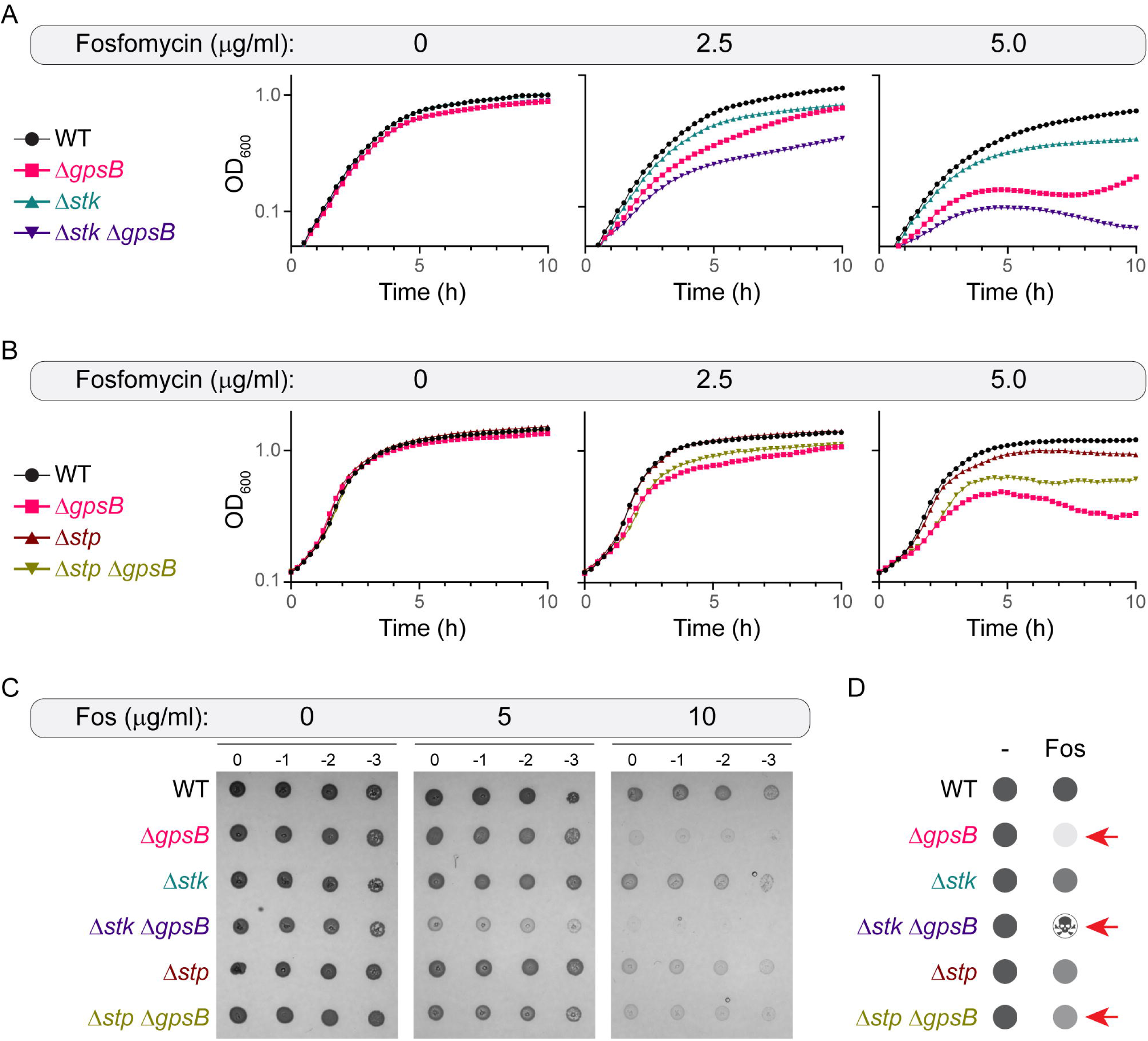
Absence of GpsB and Stk render cells hypersensitive to fosfomycin. (**A-B**) Growth curves of WT *S. aureus* (JE2), Δ*gpsB* (LM252), Δ*stk* (LM260), Δ*stk* Δ*gpsB* (LM310), Δ*stp* (LM313), and Δ*stp* Δ*gpsB* (LM314) strains grown at different indicated concentrations of fosfomycin (Fos) for 10 h at 37 °C. Shown are the representative graphs of biological replicates (n=3). (**C**) Spot titer assay of the strains listed above grown on MHA without or with Fos at indicated concentrations. Representative plate images (n=3) of overnight growth at 37 °C are shown. (**D**) Schematic summary of fosfomycin sensitivity. Red arrows point to strains lacking *gpsB* individually, or in combination with *stk* or *stp*.

In a Δ*stk* mutant strain, GpsB will be in its unphosphorylated state. Thus, we tested the opposite condition where GpsB will remain phosphorylated such as when the cognate phosphatase of Stk, Stp, is absent (**Fig. 3B**). As shown earlier, the *gpsB* deletion mutant is highly sensitive to Fos. Curiously, the strain with *stp* disruption also exhibits a mild growth defect similar to the Δ*stk* strain. This implies that Stp-dependent dephosphorylation of proteins phosphorylated by Stk is needed for optimal Fos resistance. However, in striking contrast to the Δ*stk* Δ*gpsB* strain, when compared to *gpsB* single-deletion mutant, cells lacking both *stp* and *gpsB* exhibit moderately improved growth which is more evident at 5 µg/ml Fos. This result indicates that when GpsB is absent, Stp-mediated dephosphorylation is detrimental in the context of Fos resistance. This liquid culture phenotype showing differential Fos sensitivity is also recapitulated on the solid medium (**Fig. 3C**). In this experiment also we see that the *gpsB* deletion strain exhibits heightened Fos sensitivity than the *stk* mutant (**Fig. 3D**). Similarly, Δ*stk* Δ*gpsB* is unable to grow at 10 µg/ml Fos, thus reaffirming cells lacking both Stk and GpsB are unable to handle cell wall stress. Again, as seen in liquid culture, additional deletion of *stp* alleviates severe growth defect seen in cells lacking *gpsB*. This set of data underscores the highly important role of GpsB and Stk/Stp phosphoregulation in Fos resistance (**Fig. 3D**, see arrows). Overall, our results establish that GpsB is critical for Fos resistance.

### Efficient fosfomycin resistance requires GpsB phosphoregulation

As availability of GpsB is highly critical for Fos resistance, we aimed to investigate the significance of GpsB phosphorylation. More precisely, deletion of *stk* and *stp* likely leads to global changes of *S. aureus* phosphoproteome and preclude straightforward analysis regarding the specific importance of GpsB phosphorylation. Therefore, to circumvent this, we probed the ability of GpsB phosphomutants to complement the *gpsB* null phenotype (**Fig. 4A**). As expected, unlike WT control, deletion of *gpsB* renders the cells highly sensitive to Fos. Upon complementation using a plasmid-based copy of inducible *gpsB*, we see a reversal of this phenotype. In fact, the growth of this strain reproducibly surpasses WT at both 2.5 and 5 µg/ml Fos. This implies that phospho-tunable GpsB is highly beneficial for Fos-stressed cells. Next, we tested the complementation phenotypes of T75E and T75A mutants. Intriguingly, both mutants are unable to fully complement Δ*gpsB*, when grown in the presence of Fos. However, T75A appears to be slightly better at complementing than T75E, presumably due to the presence of alternate sites for phosphoregulation (**Fig. 1**). Nevertheless, presence of either of the two phosphomutants is better than none when *gpsB* is deleted. This trend holds true on solid medium as well (**Fig. 4B**). Thus, as summarized in **Fig. 4C**, both the presence of GpsB and its phosphoregulation by Stk/Stp are needed for optimal Fos resistance. To further restrict phosphoregulation we capitalized on our triple mutants (**Fig. 1E**). Again, we see a similar trend where post-translationally tunable GpsB offering the best protection against Fos, while both AAA and EEE phosphomutants are unable to restore growth to the WT level (**Fig. S2**). In this assay also we find that the growth of phosphoablative version is modestly enhanced when compared to the phosphomimetic mutant. These results highlight the importance of GpsB and its phosphoregulation in Fos resistance.

**Fig. 4:**
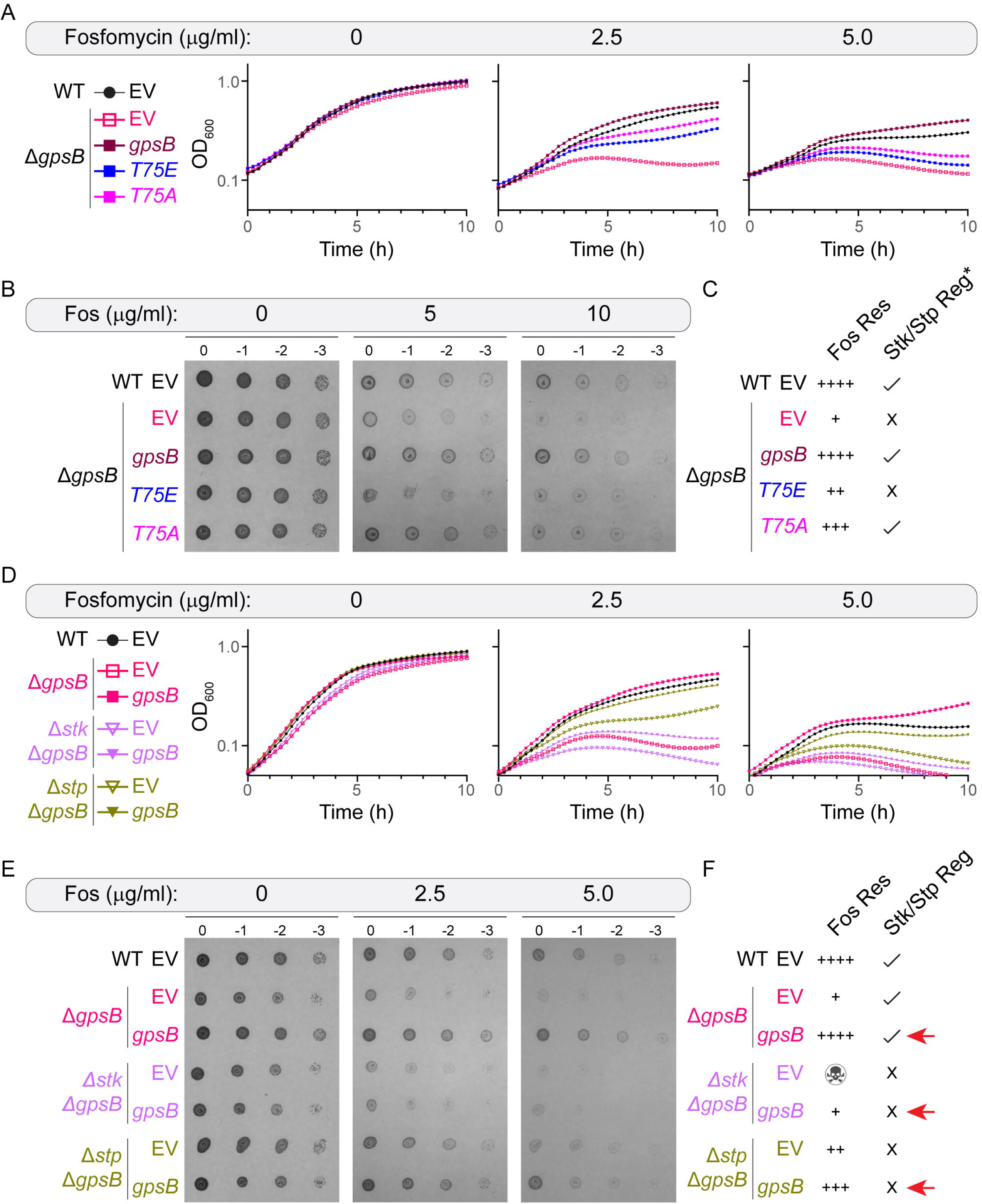
GpsB phosphoregulation is required for maximal Fos resistance. (**A-B**) Growth and spot titer assays of *S. aureus* JE2 and Δ*gpsB* strains harboring either empty vector (EV) or *gpsB*/phosphomutants. JE2 + EV (WT, LM253), Δ*gpsB* + EV (LM254), Δ*gpsB* + *gpsB* (LM255), Δ*gpsB* + T75E (LM256), and Δ*gpsB* + T75A (LM257) were grown in the absence or presence of fosfomycin (Fos) at concentrations indicated for 10 h (A) or overnight (B) at 37 °C. (**C**) Summary of the data presented in panels A and B. Fos resistance is ranked from ++++ (WT and WT-like) to + (extreme sensitivity). Presence or absence of Stk/Stp regulation (*of GpsB) is marked by and X respectively. T75A is subject to additional phosphoregulation (Fig. 1C) thus indicated as. Note: GpsB overproduction toxicity is specific to RN4220 and is not observed in JE2. (**D-E**) Liquid and solid media Fos sensitivity analyses similar to panels A and B. Strains used: JE2 + EV (WT, LM253), Δ*gpsB* + EV (LM254), Δ*gpsB* + *gpsB* (LM255), Δ*stk* Δ*gpsB* + EV (LM315), Δ*stk* Δ*gpsB* + *gpsB* (LM316), Δ*stp* Δ*gpsB* + EV (LM317), and Δ*stp* Δ*gpsB* + *gpsB* (LM318). Data representative of three independent biological replicates are shown in panels A, B, D, and E. (**F**) Schematic summary of data presented in panels D and E, similar to panel C. symbol is used to depict severe Fos sensitivity at higher concentration range. Presence or absence of Stk/Stp regulation (not just of GpsB) is marked by and X respectively. Red arrows indicate the *gpsB*-complemented strain in each genetic background to highlight that GpsB, more specifically, its phosphoregulation is imperative for optimal Fos resistance.

Next, we tested this inference independently with *stk*/*stp* deletion. Briefly, we introduced plasmid-based inducible *gpsB* in Δ*gpsB*, Δ*stk* Δ*gpsB*, and Δ*stp* Δ*gpsB* strain backgrounds and studied their Fos resistance profiles in the presence of inducer. Accordingly, GpsB produced in Δ*stk* Δ*gpsB* strain will likely be in its unphosphorylated form, while in Δ*stp* Δ*gpsB* it is amenable for phosphoregulation. Again, as shown before (**Fig. 3**), when compared to Δ*gpsB*, the strain lacking both *stk* and *gpsB* are hypersensitive to Fos while additional deletion of *stp* moderately alleviates Fos sensitivity seen in Δ*gpsB* cells (**Fig. 4D**; compare open symbols). As expected, introduction of *gpsB* fully restores Fos resistance in Δ*gpsB* strain. Surprisingly, in Δ*stk* Δ*gpsB* background, *gpsB* complementation is only mildly beneficial. This suggests that GpsB phosphorylation, in addition to its mere presence, is critical for Fos resistance. In support of this, we find that the production of plasmid-based GpsB in Δ*stp* Δ*gpsB* (where GpsB is tunable through Stk phosphorylation) restores Fos resistance to nearly WT level (**Fig. 4D**; compare filled symbols). These results are consistent when tested on solid growth medium as well (**Fig. 4E**). Taken together, our results suggest that GpsB and its post-translational regulation via Stk/Stp are key mediators of Fos resistance (**Fig. 4F**; see red arrows). Additionally, in conjunction with the data presented with phosphomutants (**Fig. 4ABC** and **Fig. S2**), we can infer that post-translational tuning of GpsB specifically is critical for achieving maximal Fos resistance.

### MurA stabilization or murA deletion reverses Fos sensitivity exhibited by the gpsB null strain

A previous report demonstrated that the enhanced sensitivity of the *stk* null strain to another cell wall targeting antibiotic, oxacillin, can be reversed by the disruption of the ReoM/ReoY pathway that is responsible for MurA turnover (18). Therefore, we wondered whether increased Fos sensitivity seen in Δ*gpsB* could also be reversed by a similar strategy. To test this possibility, we generated a *reoY* and *gpsB* double-knockout strain and tested its Fos resistance to corresponding single knockouts and WT control. As expected, we see cells lacking *gpsB* exhibit elevated sensitivity to Fos treatment (**Fig. 5A**). In contrast, we find that both the *reoY* single-deletion and the *reoY gpsB* double-deletion mutants are highly resistant to Fos, even at 10 µg/ml. The growth phenotype on Fos-containing solid medium also supports this observation (**Fig. 5B**). This result, as well as the presence of sequence weakly resembling GpsB recognition motif (R/N/K-X-X-R/K/N) (7, 13, 38), prompted us to test the interactions between MurA, MurA-paralog MurZ, ReoM, and ReoY. We used BTH assay for this purpose (**Fig. 5C**). Our results reveal that GpsB directly interacts with MurA but not MurZ (**Fig. 5F**). We also note that GpsB interacts with ReoM and ReoY. Both GpsB T75E and T75A phosphomutants also retain this interaction pattern implying that the interaction of GpsB with this set of partners is not subject to Stk-based phosphoregulation. This result constitutes the primary evidence revealing GpsB interaction with MurA, ReoM, and ReoY in *S. aureus*.

**Fig. 5:**
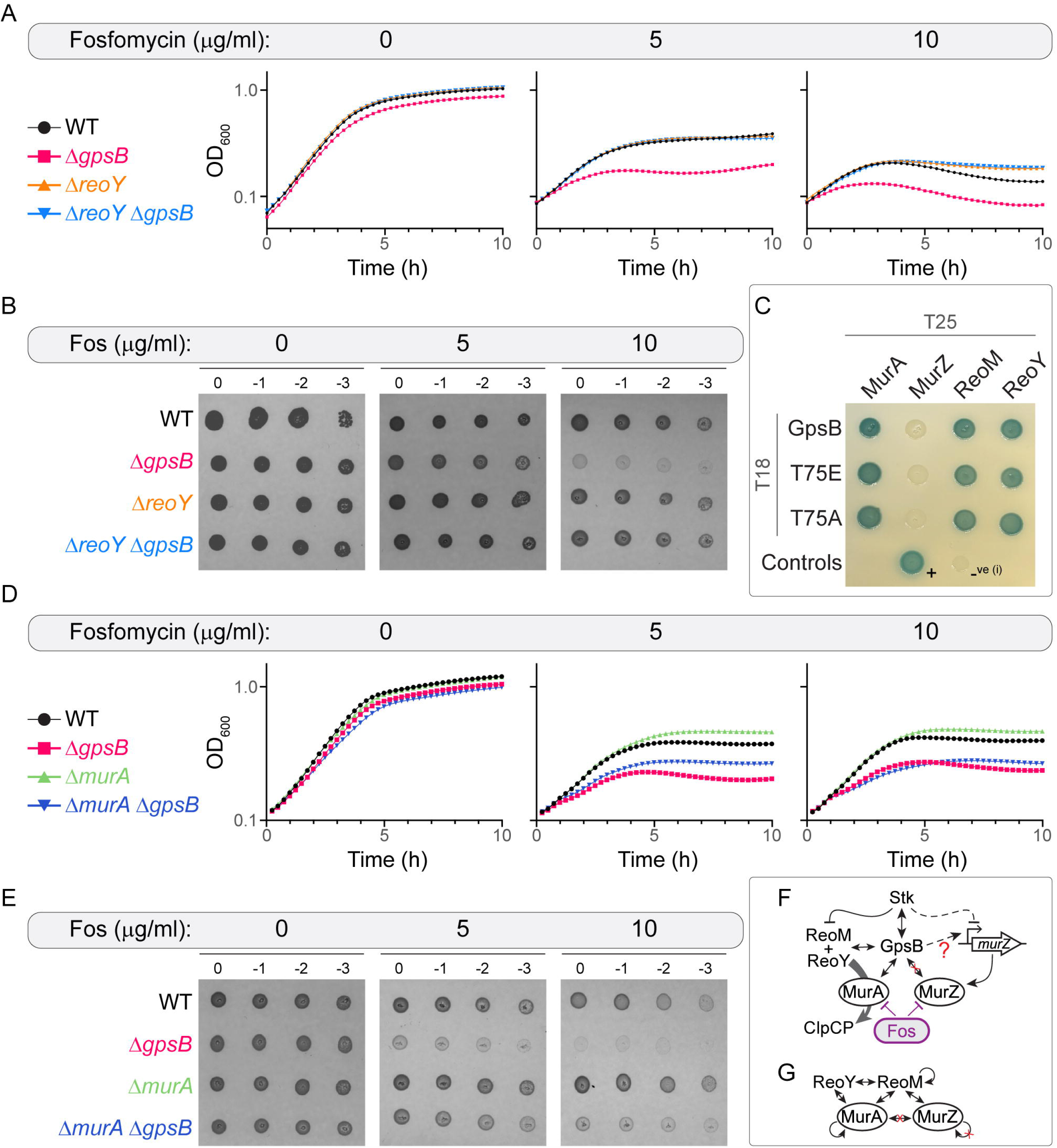
GpsB influences both MurA and MurZ pathways. (**A**) Growth curves of WT *S. aureus* (JE2), Δ*gpsB* (LM252), Δ*reoY* (LM311), Δ*reoY* Δ*gpsB* (LM312) strains grown in the absence or presence of fosfomycin (Fos) at concentrations indicated for 10 h at 37 °C. (**B**) Spot titer assay of the strains listed in panel A grown on MHA without and with fosfomycin grown overnight at 37 °C. (**C**) Bacterial two-hybrid (BTH) assay results. Interaction between GpsB or blank (-, empty vector) was tested against MurA, MurZ, ReoM, and ReoY. Additional information pertaining to BTH assay and controls are provided under methods and in the supplemental file. (**D-E**) Growth curves and spot titer assays of WT (JE2), Δ*gpsB* (LM252), Δ*murA* (LM325), Δ*murA* Δ*gpsB* (LM326) strains grown without or with Fos, as in panels A and B. (**F**) Stk-centric regulatory network related to MurA/MurZ accumulation. Fosfomycin inhibits both MurA and MurZ. Thus, increased accumulation of MurA and/or MurZ is vital for Fos resistance. The protein factors, ReoM and ReoY, together mediate the proteolytic turnover of MurA in coordination with ClpCP protease machinery. Stk-dependent phosphorylation of ReoM inhibits MurA degradation. In parallel, Stk, through phosphorylation indirectly inhibits *murZ* transcription (indicated as dashed line) by weakening the DNA binding activity of the transcription factor (VraR) responsible for *murZ* upregulation. We show that GpsB plays a central role in this regulatory pathway by directly interacting with Stk, MurA, ReoM, and ReoY. Our data suggests that GpsB may not directly interact with MurZ (red X). Yet, results presented in panels D and E implies that when MurA is absent, presence of GpsB becomes highly important to resist Fos. Therefore, we predict (red question mark; confirmed in Fig. 6) that GpsB may be involved in positively influencing *murZ* expression. (**G**) Summary of BTH assay data presented in Fig. S3A.

As MurA is the primary house-keeping enzyme targeted by Fos (29), we postulated that deletion of *murA* would phenocopy Fos sensitivity of Δ*gpsB* mutant. To test this, we repeated the Fos resistance assays with strains lacking *murA* individually or in combination with *gpsB*. In comparison to WT, and contrary to our prediction, we note that cells lacking *murA* are in fact more resistant to Fos (**Fig. 5D**). A logical explanation for this is perhaps that when MurA is absent, production of its paralog MurZ is presumably enhanced (29). To our surprise, we find that additional deletion of *gpsB* negatively affects Fos resistance in a *murA* null mutant. Interestingly, the growth of this *murA gpsB* double-deletion strain in media containing Fos is noticeably improved than *gpsB* single-deletion mutant (**Fig. 5DE**). However, this beneficial effect appears to be diminished at higher Fos concentration. This set of results reveals that: (i) deletion of *reoY*, which is expected to stabilize MurA, reverses Fos sensitivity of the *gpsB* deletion mutant (**Fig. 5AB**); and (ii) heightened Fos resistance exhibited by Δ*murA* is lost upon additional deletion of *gpsB* (**Fig. 5DE**). Together, this data underscores a previously unappreciated central function of GpsB in regulating the levels of MurA and potentially MurZ.

Stk plays an important role in MurA stabilization via ReoM phosphorylation (**Fig. 5F**). In addition, it may negatively influence *murZ* expression through VraR phosphorylation (35, 36). Thus, Stk can indirectly modulate the level of MurZ. Importantly, the antibiotic Fos inhibits the function of both MurA and MurZ (29). Therefore, stabilization of MurA and/or upregulation of *murZ* are important for Fos resistance (**Fig. 5F**). We reveal that GpsB interacts with MurA and that deletion of *reoY* reverses Fos sensitivity exhibited by the Δ*gpsB* strain, presumably due to MurA stabilization. This implies that MurA may become unstable in the absence of GpsB. Additionally, when MurA is absent, predicted upregulation of *murZ* seems to be sufficient for achieving near-WT level Fos resistance. However, importantly, this happens only in the presence of GpsB. This, in turn, suggests that MurZ stability may also be influenced by GpsB.

To further our understanding, we investigated whether MurZ is also under the control of ReoM/Y pathway we conducted a BTH assay. Excitingly, our results reveal that MurZ and ReoM are indeed interaction partners (**Fig. S3A** and **Fig. 5G**). We also find that ReoM and MurA self-interacts, which is not the case with MurZ. This supports the notion that MurZ likely functions as a monomer (34). Additionally, we do not see a direct MurA-MurZ interaction in our assay. However, we see that MurZ directly interacts with ReoM (42, 46), which may bridge the interaction MurA and MurZ (47). Although we do not observe MurZ-ReoY interaction, it is plausible that MurZ stability and/or its enzymatic function is directly influenced by ReoM (34, 46, 48). However, as GpsB does not directly interact with MurZ and lack of MurA is not detrimental in the context of Fos resistance, we predict that GpsB may influence *murZ* upregulation instead. Taken together, our data implies that GpsB, either directly or indirectly, is critical for optimal accumulation of both MurA and MurZ.

### murZ expression is compromised in the absence of GpsB

It has been demonstrated that *murZ* is upregulated in response to cell wall stress, while *murA* is not (29, 49). Additionally, our data reveals that MurZ pathway is linked to GpsB (**Fig. 5F**). Thus, we sought to investigate the *murZ* promoter activity. For this purpose, we generated a luciferase-based *murZ* (*P_murZ_-lux*) transcriptional reporter (50). We then introduced this reporter in WT, Δ*gpsB*, Δ*stk*, Δ*stk* Δ*gpsB*, Δ*murA*, and Δ*murA* Δ*gpsB* strain backgrounds. Fos sensitivity of these strains as monitored by growth on plate (**Fig. 6A**) and OD_600_ (**Fig. S3BC**) resemble those without the reporter plasmid. Imaging these plates via luminescence channel revealed that the *P_murZ_* activity in *gpsB* null strain is markedly lower than WT control even in the absence of Fos (**Fig. 6B**). The reduction of the luminescence signal in Δ*gpsB* is more pronounced in the presence of Fos, which is also evident in liquid culture assay (**Fig. S3BC**, see bottom panels). Curiously, in WT, we see increased *murZ* promoter activity during logarithmic growth phase even in untreated condition. This suggests that MurZ is also utilized by non-antibiotic stressed cells presumably to support efficient growth rate.

**Fig 6:**
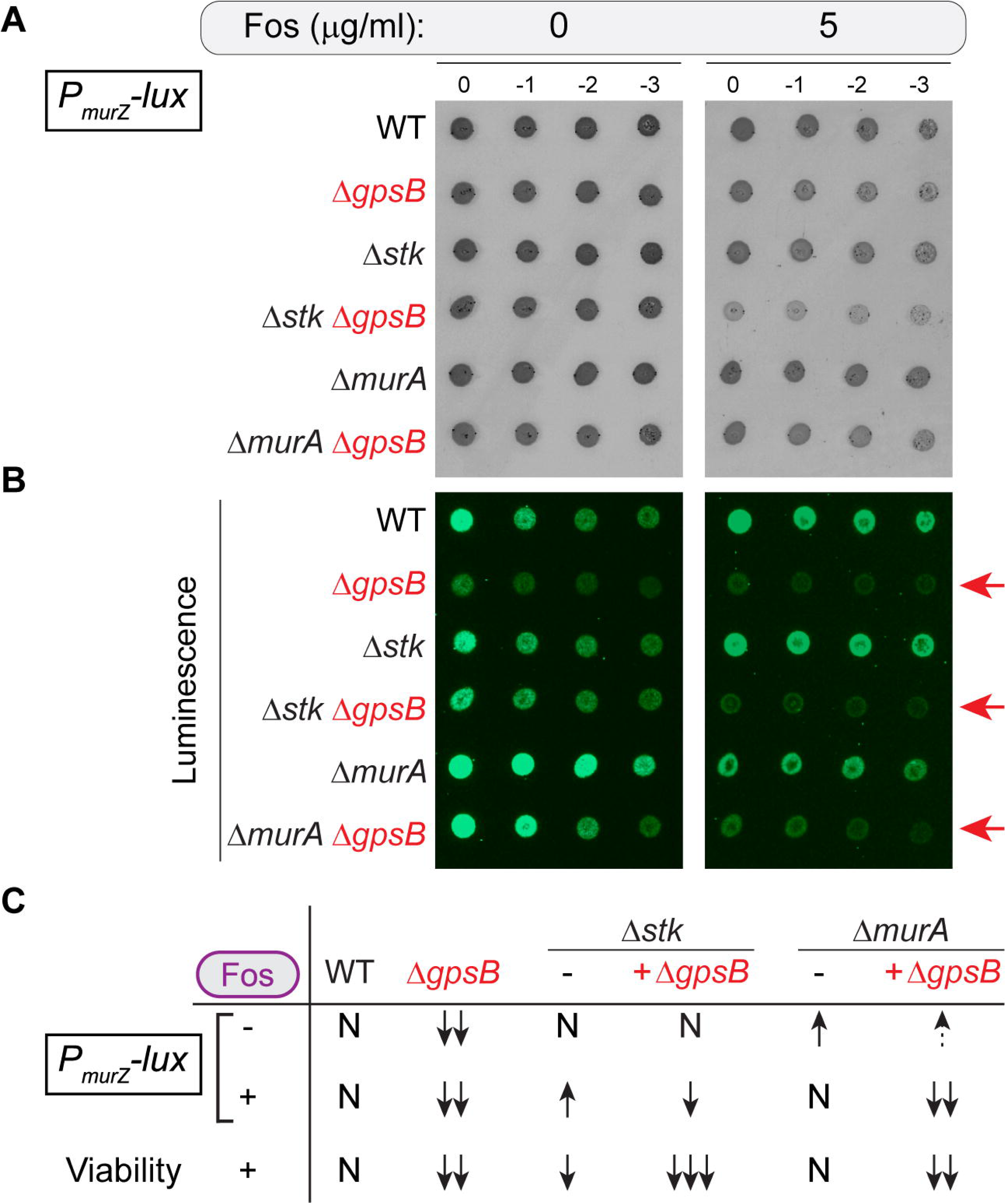
Presence of GpsB is critical for *murZ* upregulation. (**A-B**) Spot titer assay of strains (listed below) harboring *P_murZ_-lux* transcriptional reporter grown on MHA without (left panels) or with 5 µg/ml fosfomycin (right panels) grown overnight at 37 °C. Reporter strains used: WT JE2 (LM329), Δ*gpsB* (LM330), Δ*stk* (LM331), Δ*stk* Δ*gpsB* (LM332), Δ*murA* (LM333), and Δ*murA* Δ*gpsB* (LM334). Representative images of plates imaged through transillumination (panel A) or chemiluminescence (panel B) are shown, n=3. Red arrows point to strains in which *gpsB* is deleted individually or together with *stk* or *murA* to emphasize diminished *murZ* promoter activity. (**C**) Summary of the data presented in panels A, B, and Fig. S3BC. Arrows (↑) or (↓) indicate increased or decreased *P_murZ_-lux* activity or viability relative to WT (N, normal). Weak increase (dashed ↑) and strong decrease (increasing number of ↓) are symbolized to aid in the interpretation of the results.

As we find that the *gpsB* null strain is more sensitive to Fos than the Δ*stk* strain (**Fig. 3A**), we monitored the *P_murZ_* activity in this background. As noted previously, cells lacking *stk* alone exhibits only mild growth inhibition on Fos, while the growth of Δ*stk* Δ*gpsB* is severely hindered. In support of our prediction, Δ*stk* cells exhibit higher luminescence than the WT control in liquid culture under both untreated and, more obviously, in Fos-treated conditions (**Fig. S3B**). Thus, increased expression of *murZ* possibly leads to minimal loss of Fos resistance seen in the *stk* null strain. Intriguingly, however, cells lacking both *stk* and *gpsB* produce higher luminescence signal when compared to the *gpsB* deletion strain, even in untreated conditions in the absence of any growth deficiency in both solid and liquid media. Why even with higher *murZ* promoter activity than Δ*gpsB*, the Δ*stk* Δ*gpsB* double-deletion mutant displays heightened Fos sensitivity is puzzling (**Fig. S3B**). A potential possibility is that perhaps MurZ stability/catalytic function, in addition to its expression, is also negatively affected in the absence of GpsB, presumably through ReoM interaction (**Fig. 5G**). Hence, in the absence of both GpsB and Stk, increased expression of *murZ* is insufficient to support growth in the presence of Fos. However, this possibility remains to be tested.

In the context of Δ*murA*, as expected, the *P_murZ_-lux* reporter activity is elevated in the absence of Fos (**Fig. 6B**), but closely resembles WT in the presence of Fos as seen on both solid and liquid media (**Fig. S3C**, see bottom right panel). This implies that in the presence of additional regulators such as Stk and GpsB, WT-level *murZ* expression is sufficient to confer maximal Fos resistance (**Fig. 5D**). In the Δ*murA* Δ*gpsB* strain background, the *P_murZ_* activity mimics, albeit somewhat weakly, the trend of the *murA* null mutant in the absence of Fos. However, upon Fos treatment, the *murZ* expression profile of this strain is highly impaired similar to the *gpsB* single-deletion mutant in both solid (**Fig. 6B**) and liquid (**Fig. S3C**, compare bottom two panels) media conditions. It is worth noting that in the absence of *P_murZ_-lux* reporter plasmid (**Fig. 5E**), we observe modest improvement in Fos resistance for Δ*murA* Δ*gpsB* when compared to Δ*gpsB*. Therefore, it is safe to infer that increased expression of *murZ* is responsible for improved Fos resistance in the *murA gpsB* double-deletion at lower antibiotic concentration. In contrast, at higher Fos concentrations this protective effect is absent. This implies that, at higher Fos concentration, the subdued *murZ* promoter activity in Δ*murA* Δ*gpsB* is likely responsible for increased Fos sensitivity (**Fig. 5DE**). From these results, we deduce that in the absence of GpsB, stress-responsive *murZ* upregulation is compromised which render cells lacking *gpsB* highly susceptible to Fos. Thus, as summarized in **Fig. 6C**, both *murZ* expression and the viability of all mutants lacking *gpsB* are compromised in the presence of Fos. Overall, this dataset further strengthens the notion that GpsB is critical for reinforcing the MurZ pathway.

#### GpsB collaborates with VraRS to provide Fos resistance

As the cell wall stress responsive VraSR TCS mediates *murZ* upregulation (35, 51), we next probed its link to GpsB. We hypothesized that deleting *vraR*/*S* may phenocopy Δ*gpsB* due to impaired *murZ* expression. Contrary to our prediction, as shown in **Fig. 7A**, we find that cells lacking either *vraR* or *vraS* are more sensitive to Fos than the *gpsB* null mutant. However, when both GpsB and VraR/S are absent, cells become hypersensitive to Fos (**Fig. 7A**, see arrowheads). Our liquid culture data is also consistent with this observation (**Fig. S4AB**). Therefore, it appears that GpsB influences the VraSR TCS pathway. To test whether protein-protein interaction underlie this effect, we tested the interaction between GpsB and VraR/S via BTH assay. Remarkably, our results reveal that GpsB directly interacts with VraR (**Fig. 7B**). Additionally, we see that the T75E phosphomimetic version of GpsB, but not T75A, interacts with both VraR and VraS. This is the first known direct connection between GpsB and TCS system to our knowledge.

**Fig 7:**
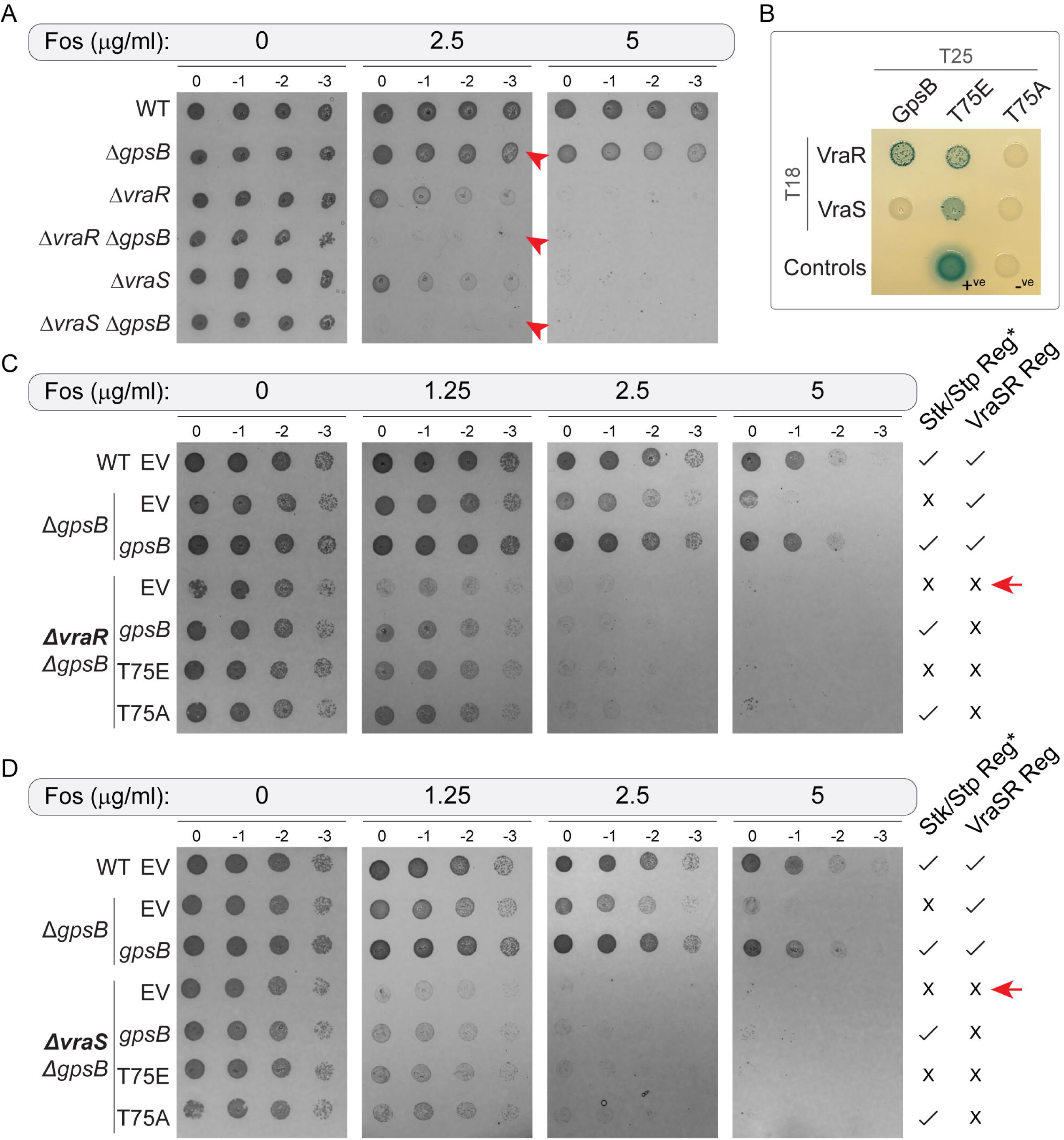
GpsB collaborates with the cell wall stress sensing VraSR two-component system. **(A)** Spot titer assay of WT *S. aureus* (JE2), Δ*gpsB* (LM252), Δ*vraR* (LM338), Δ*vraR* Δ*gpsB* (LM339), Δ*vraS* (LM336), Δ*vraS* Δ*gpsB* (LM337) strains grown on MHA without and with fosfomycin at concentrations indicated. Red arrowheads indicate strains lacking *gpsB*. **(B)** Bacterial two-hybrid (BTH) assay results. Interaction between GpsB and its phosphomutant alleles were tested against VraR and VraS. Information regarding positive and negative controls are provided in the methods section. **(C-D)** Spot titer assays of *S. aureus* JE2, Δ*gpsB* and Δ*vraR* Δ*gpsB* (C) Δ*vraS* Δ*gpsB* (D) harboring either empty vector (EV) or *gpsB*/phosphomutants grown in the absence or presence of fosfomycin (Fos) at indicated concentrations. Strains used: JE2 + EV (WT, LM253), Δ*gpsB* + EV (LM254), Δ*gpsB* + *gpsB* (LM255), Δ*vraR* Δ*gpsB* + EV (LM344), Δ*vraR* Δ*gpsB* + gpsB (LM345), Δ*vraR* Δ*gpsB* + T75E (LM346), Δ*vraR* Δ*gpsB* + T75A (LM347), Δ*vraS* Δ*gpsB* + EV (LM340), Δ*vraS* Δ*gpsB* + *gpsB* (LM341), Δ*vraS* Δ*gpsB* + T75E (LM342), and Δ*vraS* Δ*gpsB* + T75A (LM343) were overnight at 37 °C. Images representative of plates grown overnight at 37 °C are shown (n=3). Presence or absence of regulation by Stk/Stp (*of GpsB) and VraSR TCS are marked by and X respectively. As T75A may be available for additional phosphoregulation (Fig. 1C), it is indicated as. Red arrows indicate strains with the highest Fos sensitivity.

As we uncovered a tripartite stress response network involving GpsB, Stk, and VraSR, we aimed to further dissect their respective roles. For this purpose, we generated a *gpsB* and *vraR*/*S* double-deletion strain and complemented with GpsB or T75E/A phosphomutants (**Fig. 7CD**). First, we confirmed the Fos sensitivity of Δ*gpsB* is fully complemented by *gpsB* as shown before (**Fig. 4B**). We see that both Δ*vraR* Δ*gpsB* (**Fig. 7C** and **Fig. S4C**) and Δ*vraS* Δ*gpsB* (**Fig. 7D** and **Fig. S4D**) exhibit heightened Fos sensitivity (see 1.25 µg/ml panels). Complementation of *gpsB* or its phosphomutants moderately improves growth at 1.25 µg/ml Fos concentration in both solid and liquid media. Curiously, however, they are insufficient to support growth at higher concentration range. Therefore, our results suggest that intact VraSR TCS is critical for optimal Fos resistance (Fig. 7CD, see red arrows). Collectively, our results shed light on the previously unknown coordination between GpsB and the VraSR two-component cell wall stress response system.

## Discussion

The various partners and functions of GpsB is relatively well characterized in several organisms including *S. aureus* (5–9, 14, 15). Consequently, elucidating the role of GpsB in antibiotic resistance is an active area of investigation (19, 33, 52–54). In this study, we sought to reveal the physiological significance of GpsB phosphorylation and its potential role in antibiotic resistance in *S. aureus*. The results shown in **Fig. 1** revealed that although GpsB is phosphorylated at multiple residues, chief among them are three specific phosphosites (T21, T75, and S76) which appear to be individually sufficient for phosphoregulation. However, as our readout was solely based on overproduction toxicity specifically in *S. aureus* RN4220 background, it remains possible that the other sites are used for additional Stk-mediated fine-tuning of GpsB that we did not capture in our experimental conditions. Using T75E phosphomimetic mutation as proxy for phosphorylation, we observed that GpsB phosphostatus toggles GpsB partner preference (**Fig. 2**). Specifically, our data revealed that the T75E version of GpsB strongly associates with Stk. Remarkably, similar observation was reported in *Streptococcus pneumoniae* as well (21). On the contrary, the interaction between GpsB and PBP2 is favored when the T75 residue of the former is unmodified. Equipped with this new knowledge, we then probed the role of GpsB and its phosphoregulation in antibiotic resistance.

As Stk signaling is linked to cell wall homeostasis (26), we investigated the importance of GpsB phosphoregulation in the context of fosfomycin antibiotic which inhibits MurA and MurZ to deplete the availability of peptidoglycan precursors and induces cell envelope stress (29). While deletion of *stk* resulted in a modest decrease in Fos resistance, to our surprise, we observed that cells lacking *gpsB* are highly sensitive to Fos (**Fig. 3D**). Intriguingly, deletion of *stk* and *gpsB* further worsened Fos sensitivity. In contrast, deletion of both *stp* and *gpsB* alleviated the Fos sensitivity exhibited by cells lacking *gpsB* alone. Thus, in the absence of GpsB∼P interaction/stimulation of Stk (**Fig. 2E**), removal of Stp presumably elevates the levels of substrate proteins phosphorylated by Stk, such as ReoM (discussed below), which is helpful in the presence of Fos. Notably, GpsB-dependent stimulation of Stk kinase activity is experimentally confirmed in other organisms (20, 21, 33, 55). Furthermore, using complementation experiments, we discovered that only the phospho-tunable version of GpsB can provide maximal protection against Fos (**Fig. 4C** and **Fig. S2**). Curiously, however, the mere presence of GpsB does confer moderate Fos resistance specifically when Stp, but not Stk, is absent. This suggests that phosphorylation of at least some subpopulation of GpsB is important for reversing Fos sensitivity **(Fig. 4F**).

MurA turnover is mediated by the ReoM/Y complex which serves as the ClpCP protease adapter in multiple organisms (18, 30, 42, 56). Relevant to this study, phosphorylation of ReoM by S/T kinase disrupts this pathway and allows MurA to accumulate. Accordingly, the heightened antibiotic sensitivity exhibited by the *stk* null mutant in *S. aureus* is reversed by the deletion of *clpP*, *reoM*, or *reoY* (18). This observation established that this Stk/MurA link is conserved in S. aureus as well (**Fig. 5F**). We saw that the increased Fos sensitivity seen in cells lacking *gpsB* was also reversed by *reoY* deletion (**Fig. 5AB**). This finding hinted that, in the absence of GpsB, MurA level is presumably depleted. We also uncovered that GpsB directly interacts with ReoM, ReoY, and MurA, but not MurZ (**Fig. 5C**). Thus, we believe GpsB may support MurA accumulation independent of Stk-ReoM pathway (**Fig. 5F**). This would explain why Fos-treated cells fare well as long as GpsB is present even in the absence of Stk/Stp regulation. Next, we pondered whether abolishing the MurA pathway depicted in **Fig. 5F** would result in heightened susceptibility to Fos. Remarkably, that is not the case as MurA paralog, MurZ, seems to be sufficient for providing WT-level Fos resistance (**Fig. 5DE**). However, Fos resistance seen in Δ*murA* was diminished in the absence of GpsB. Thus, GpsB is central to both MurA and MurZ pathway and this role of GpsB becomes critical in the presence of fosfomycin (**Fig. 5F**).

As our results failed to show a direct interaction with MurZ, we wondered whether stress-adaptive *murZ* promoter activity relies on GpsB. The VraRS two-component cell wall stress signal sensing system is responsible for *murZ* upregulation (36). Importantly, the DNA binding activity of VraR is shown to be impaired upon Stk phosphorylation (35). Thus, *stk* deletion should alleviate that negative regulation and lead to increased expression of *murZ*. In fact, that is indeed what we observed (**Fig. 6C**). Similarly, as expected, we saw WT-like *murZ* expression pattern in the mutant lacking *murA*. Astoundingly, regardless of whether GpsB is individually absent or in combination with Stk or MurA, *murZ* expression level is significantly reduced especially in the presence of Fos. Even when *murZ* transcription is higher in the strain lacking both *stk* and *gpsB* than in the Δ*gpsB* mutant, the double-deletion strain exhibits heightened sensitivity to Fos (**Fig. 6C**). This suggested that the presence of GpsB might offer some protection to MurZ perhaps through ReoM sequestration (**Fig. 5FG**). Additionally in strains lacking *gpsB*, even in the absence of Fos and any growth defects, the *murZ* promoter reporter activity showed marked decrease. This prompted us to investigate the potential link between GpsB and VraSR TCS. While the absence of VraS/R increases Fos sensitivity, our results showed that this is worsened by additional deletion of *gpsB* (**Fig. 7A**). Our BTH analysis uncovered the interaction between GpsB and VraR and potential phosphorylation-dependent interaction between GpsB and VraS (**Fig. 7B**). Furthermore, our data elucidated that the VraSR system, reinforced by GpsB and Stk, plays a predominant role in providing Fos resistance. In accordance with of our observation, another recent report also found similar sensitivity for mutants lacking *vraSR* and *gpsB* to a highly clinically-relevant antibiotic, dalbavancin (53). Thus, it appears that the antibiotic resistance-conferring collaboration between GpsB and VraSR may extend beyond Fos to other cell envelope targeting antibiotics.

Our results highlight the central role played by GpsB in achieving cell wall homeostasis (**Fig. 8A**). Through direct interaction with partner proteins such as PBP2, GpsB monitors cell wall synthesis. Accordingly, GpsB may be able to perceive peptidoglycan synthesis impairment. Alternatively, Stk may directly sense cell envelope stress independent of GpsB depending on the type of stress (54). As our data suggests that GpsB phosphorylation likely favors stronger interaction with Stk, this would serve to further amplify the stress signal. In turn, Stk stabilizes MurA through ReoM phosphorylation, which interferes with the ReoY/ClpCP-mediated proteolysis. As inhibition of MurA proteolysis is presumably the quickest and most energy efficient way to alleviate cell wall stress, Stk phosphorylation of VraR restrains *murZ* upregulation. Consistently, we observe increased *murZ* promoter activity when Stk is absent in the presence of Fos. Intriguingly, *murZ* upregulation is diminished when *gpsB* is deleted. This strengthens the notion that GpsB positively influences *murZ* expression through VraSR system independent of Stk. Additionally, we also detect direct interactions between GpsB-MurA, GpsB-ReoM, as well as ReoM-MurA and ReoM-MurZ. Thus, it is plausible GpsB, either directly or indirectly through alleviating ReoM binding, stabilizes MurA and MurZ.

**Fig 8:**
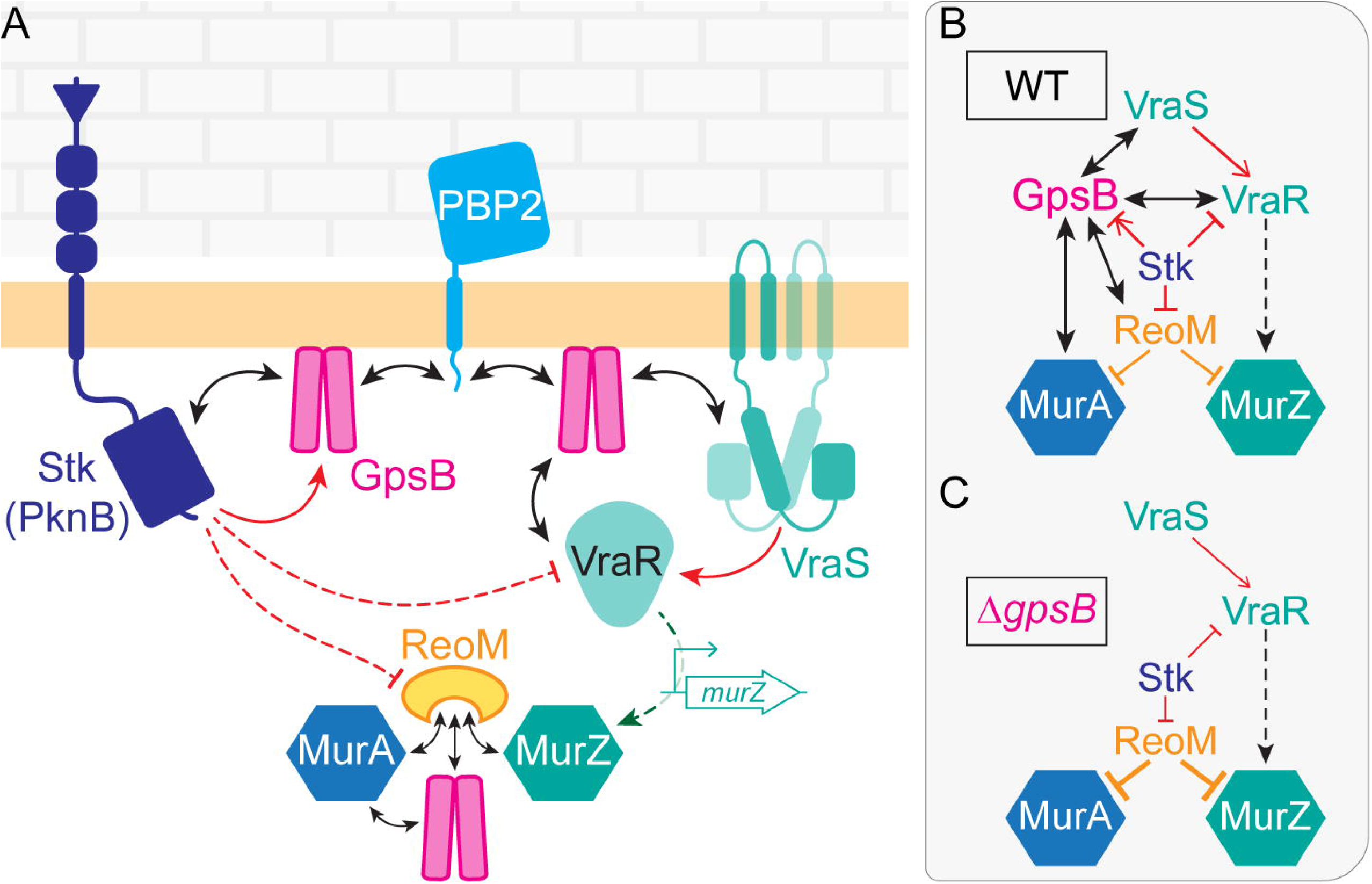
GpsB-Stk-VraSR form a cell wall stress surveillance network to maintain cell envelope integrity. (**A**) Schematic model to illustrate the pivotal role of GpsB in regulating cell wall synthesis. We presume that GpsB-PBP2 interaction, and that of other partners associated with cell wall synthesis such as PBP4/PBP2a, is weakened during cell wall stress. Additionally, cell envelope stress is directly sensed by Stk through its extracellular PASTA domains. In either scenario, GpsB becomes more available for phosphorylation by Stk. This, in turn, strengthens GpsB-Stk interaction and leads to the stimulation of Stk kinase activity. Thus, GpsB-Stk interaction serves as a “signal amplifier” to broadcast cell envelope stress. Consequently, Stk phosphorylates ReoM and VraR to inhibit MurA turnover (via ReoY/ClpCP, not shown for simplicity) and *murZ* upregulation, respectively. We show that GpsB directly interacts with MurA and ReoM, but not MurZ (Fig. 5FG). Similarly, we also reveal that both MurA and MurZ interact with ReoM. Moreover, we find that GpsB is critical for optimal *murZ* expression (Fig. 6C) presumably through VraSR (Fig. 7B). Direct interaction between proteins are depicted as reversible arrows (↔). S/T and H/D phosphorylation by Stk and VraSR respectively are indicated by red arrows. (**B-C**) Explanation for Fos resistance in wild type (B) and heightened sensitivity in the *gpsB* null mutant (C). It is conceivable, based on the model provided in panel A, that the coordination between GpsB/Stk/VraSR is crucial for enhanced Fos resistance seen in WT. Briefly, VraSR TCS transcriptionally controls the production of MurZ. ReoM negatively regulates MurA/Z through either facilitating their proteolytic turnover and/or potentially inhibiting their enzymatic function. GpsB by directly interacting with its partners Stk, VraS/R, MurA, and ReoM/Y helps optimize the accumulation of MurA and production of MurZ. Centrally-positioned Stk, via phosphorylation, toggles the partners of GpsB, relieves ReoM inhibition of MurA/Z, and restrains VraR-dependent upregulation of *murZ*. In contrast, cell lacking *gpsB* are unable to properly tune the accumulation/function of MurA/Z. Therefore, when GpsB is absent, *S. aureus* cells are unable to mount a coordinated response required to address cell wall stress.

With the help of this model, we can explain the various Fos sensitivity phenotypes we report in this article (**Fig. 8B**). The WT cells employ the interconnected cell wall stress surveillance network involving GpsB, Stk, VraSR pathways to harmonize the need-based availability of peptidoglycan precursors. Conversely, the heightened Fos sensitivity observed in the *gpsB* deletion mutant is due to dysfunctional Stk kinase activity, loss of stable MurA, lack of optimal *murZ* upregulation, and possibly inhibition of MurZ function by ReoM (**Fig. 8C**). Thus, additional deletion of *reoY* in the strain lacking *gpsB* confers increased Fos resistance (**Fig. 5AB**). This model reveals why the deletion of *stk* alone only modestly impacts Fos resistance (**Fig. 3**) and *murA* deletion by itself is well tolerated even in the presence of Fos (**Fig. 5DE**). However, why deletion of both *murA* and *gpsB* improves the Fos resistance compared to the *gpsB* single-deletion strain, albeit only at low concentration range, is intriguing. In this case, ReoM is phosphorylated by Stk, perhaps weakly due to the lack of GpsB mediated stimulation. This, in turn, relieves MurZ-ReoM interaction/inhibition as recently proposed in other organisms (46, 48). Additionally, the DNA-binding function of VraR is not sufficiently affected by the less-active Stk and this leads to elevated *murZ* expression as evidenced in the absence of Fos (**Fig. 6BC**). As this maybe the case in cells lacking *gpsB* as well, better growth of the Δ*murA* Δ*gpsB* strain at low concentrations of Fos alludes to the possibility that the MurA-ReoM-MurZ complex is likely detrimental for MurZ accumulation or function (46). In support of this, we find that *murA* deletion is indeed beneficial (**Fig. 5DE**). However, at high Fos concentration, presence of GpsB (and Stk) becomes critical because VraR also upregulates cell wall synthesis associated genes such as PBP2 (36, 57). Conceivably, this would create increased demand for cell wall precursors and strain the system especially when the cells are solely relying on MurZ. In summary, our results reveal the central role that GpsB plays in maintaining *S. aureus* cell envelope integrity and mediating antibiotic resistance.

## Materials and Methods

### Bacterial strains and growth conditions

*S. aureus* strains were derived from methicillin-resistant USA300 LAC derivative JE2 (58), or from the restriction-deficient cloning intermediate RN4220 (59). *gpsB::kan* mutant is a kind gift from Dr. Simon Foster (14). Chromosomal gene knockout strains (*stk*::*tn*, *stp*::*tn*, *murA*::*tn*, *reoY*::*tn*) were obtained from the Nebraska Transposon Mutant Library (NTML) (58), freshly transduced into relevant strain backgrounds, and specific gene disruptions were confirmed by PCR. Double mutant combinations (e.g., Δ*stk* Δ*gpsB*, Δ*murA* Δ*gpsB*, Δ*reoY* Δ*gpsB*) were constructed by transducing the relevant resistance marker-linked allele into the appropriate single-mutant background. Selection was conducted appropriate antibiotic containing plates and the corresponding mutations were PCR verified. All strains and oligonucleotides used in this study are listed in Tables S1 and S2 respectively. *E. coli* strains DH5α, BL21-DE3, and BTH101 were used for cloning, protein purification, and bacterial two-hybrid analysis respectively, as described below. *S. aureus* was cultured in tryptic soy broth (TSB), tryptic soy agar (TSA), Mueller-Hinton broth (MHB), or Mueller-Hinton agar (MHA). *E. coli* in lysogeny broth (LB) or on LB agar. Media were supplemented with 10 µg/ml chloramphenicol (*S. aureus*, plasmid maintenance of pCL15 (60) or pXen1 (50) plasmids) or 100 µg/mL ampicillin and 50 µg/mL kanamycin (*E. coli*, BTH plasmids (43)) as needed. Expression from the IPTG-inducible *P_spac_* promoter on pCL15 (60) was induced with 1 mM IPTG.

### GpsB overproduction toxicity assay

Overnight cultures of RN4220 Δ*gpsB* harboring the empty vector or the indicated IPTG-inducible *gpsB* allele were standardized to OD_600_ = 0.1, grown to mid-log and serially diluted tenfold in TSB. Then, 1 µl of each dilution was spotted onto TSA containing 10 µg/ml chloramphenicol, either without inducer or with 1 mM IPTG, following the spot titer format previously described for *S. aureus* GpsB overproduction assays (6). Plates were imaged after overnight incubation at 37 °C. Assays were performed in 3 biological replicates (x 3 technical replicates); representative plates pictures are shown.

### Immunoblot analysis

Overnight cultures of the indicated strains were diluted 1:100 into fresh TSB containing 10 µg/mL chloramphenicol, grown at 37 °C to mid-logarithmic phase (OD_600_ ∼0.5), and treated with IPTG (1 mM) for 3 h. Cells were harvested, resuspended in PBS with 5 µl lysostaphin (1 mg/ml in 20 mM sodium acetate), and incubated for 30 min at 37 °C, following the lysis method described previously (6). Lysates were separated by SDS-PAGE, transferred, and probed with anti-GpsB or anti-SigA antibodies (5). Representative blots from 3 biological replicates are shown.

### Bacterial two-hybrid assay

Bacterial two-hybrid (BTH) analysis (43), was performed as described previously (6). Plasmid pairs encoding T18 and T25 adenylate cyclase fragment fusions were co-transformed into *E. coli* BTH101 (*cya^−^*) and selected on LB agar containing 100 µg/ml ampicillin and 50 µg/ml kanamycin at 30 °C. Individual co-transformants were grown overnight in LB with both antibiotics at 30°C, then spotted onto LB agar containing ampicillin (100 µg/ml), kanamycin (50 µg/ml), X-gal (40 µg/ml) and IPTG (1 mM) to investigate protein-protein interaction. Positive and negative controls: co-transformation of pUT18-*zip* with pUT25-*zip* served as the positive control (+ve; blue color change) (43); empty pEB354 with empty pEB355 served as the negative control (-ve^(i)^), as well as empty pEB354 with pUT18-GpsB (-ve^(ii)^). A flexible linker was appended to GpsB constructs (61, 62). All interactions were assessed by a minimum of 3 independent biological replicates. Representative plates pictures are shown.

### FtsZ/GpsB in vitro assays

FtsZ and GpsB purification procedures were carried out as described previously (13). FtsZ-GpsB binding assay: This was performed using fluorescein 5(6)-isothiocyanate (FITC) labeled FtsZ as described previously (13). Briefly, FITC-FtsZ (100 nM) was incubated with 20 µM of GpsB or GpsB-T75E in 25 mM HEPES (pH 6.7) for 10 min at 25 °C. The fluorescence spectra of FITC-FtsZ (510–650 nm) were monitored by exciting the reaction mixtures at 495 nm. The maximum fluorescence (ΔF_max_) observed at 523 nm was calculated with FITC-FtsZ and subtracted by FITC-FtsZ with GpsB or GpsB-T75E. The change in maximum fluorescence was calculated relative to FITC-FtsZ alone or with the addition of GpsB. The data plotted are the result of three independent experiments. Statistical significance was assessed by Welch’s t-test. GTPase assay: The effect of GpsB or GpsB-T75E on stimulating the GTPase activity of FtsZ was determined via a malachite green assay (Sigma). Briefly, FtsZ (30 µM) was incubated with GpsB (WT or T75E) or alone in 25 mM HEPES buffer containing 140 mM KCl, 5 mM MgCl_2_ and 2 mM GTP at 37 °C for 15 min. Then, reaction mixtures were incubated with the working reagent provided with the kit for an additional 30 min in the dark. A 100 ul aliquot of each reaction was then placed in a black 96-well plate. Inorganic phosphate release was then determined by measuring the corresponding absorbance at 650 nm in a microplate reader (BioTeK Synergy H1). The amount of free phosphate released was determined using a standard curve using the phosphate standard provided with the kit. Statistical significance was assessed by Welch’s t-test.

### Antibiotic sensitivity assays in liquid culture

Overnight cultures were diluted to OD_600_ = 0.1 in fresh MHB supplemented, as needed, with 10 µg/ml chloramphenicol for plasmid maintenance and 1 mM IPTG for complementation strains. Then, 100 µl aliquots were dispensed into 96-well plates. Fosfomycin was diluted 2-fold down the plate to the desired concentration using multichannel pipette. OD_600_ was recorded in a microplate reader (BioTeK Synergy H1) every 15 min for 10 h at 37 °C with shaking. Each experiment was performed with 3 biological replicates (x 3 technical replicates). Representative growth curves are shown.

### Antibiotic sensitivity assays on solid medium

Overnight cultures were normalized to OD_600_ = 0.1 in fresh MHB, grown to mid-logarithmic phase (OD_600_ ∼0.5) and serially diluted tenfold in MHB to 10^-3^. Then, 1 µl of each dilution was spotted onto MHA plate containing the indicated concentration of fosfomycin (additionally containing 10 µg/mL chloramphenicol and 1 mM IPTG as needed). Plates were imaged after overnight incubation at 37 °C; representative plates pictures are shown. Assays were performed in 3 biological replicates.

### murZ transcriptional reporter assays

Liquid and solid-medium *P_murZ_-lux* assays were performed as described above (antibiotic sensitivity assays), with the addition of luminescence measurement (liquid: every 15 min alongside OD_600_; solid medium: imaged under chemiluminescence with no external illumination). Assays were performed in 3 biological replicates. Representative growth curves and plates pictures are shown.

### Bioinformatics

Multiple sequence alignment and secondary structure features of GpsB sequences from *S. aureus* NCTC8325, *B. subtilis* 168, *L. monocytogenes* 10403S, *S. pneumoniae* D39, and *E. faecalis* V583 were obtained using Clustal Omega and Jalview (63, 64). Additional annotations were made to highlight phosphosites and represent details derived from solved structures and available literature.

## Acknowledgements

We thank Dr. Simon Foster for strains. We appreciate the Eswara laboratory members for their valuable input, thoughtful critiques, and engaging discussions throughout the course of this study. This work was supported by the National Institutes of Health (R35GM133617 to PJE) and Center for Antimicrobial Resistance award from the University of South Florida (PJE). The funders had no role in study design, data collection and analysis, decision to publish, or preparation of the manuscript.

## Author Contributions

Study design (LJM and PJE), strain construction and data acquisition (LJM, OP, AK, JC, BB, and LRH), data analysis (LJM, AK, and LRH), manuscript writing (LJM and PJE), manuscript comments (AK and LRH), resources (PJE), funding (PJE), and supervision (PJE).

## References

1. Jesudason T. WHO publishes updated list of bacterial priority pathogens. Lancet Microbe. 2024;5(9):100940.

2. Kariuki S. Global burden of antimicrobial resistance and forecasts to 2050. Lancet. 2024;404(10459):1172–3.

3. David MZ, Daum RS. Treatment of Staphylococcus aureus Infections. Curr Top Microbiol Immunol. 2017;409:325–83.

4. Tong SYC, Fowler VG, Jr., Skalla L, Holland TL. Management of Staphylococcus aureus Bacteremia: A Review. JAMA. 2025;334(9):798–808.

5. Eswara PJ, Brzozowski RS, Viola MG, Graham G, Spanoudis C, Trebino C, et al. An essential *Staphylococcus aureus* cell division protein directly regulates FtsZ dynamics. Elife. 2018;7.

6. Hammond LR, Sacco MD, Khan SJ, Spanoudis C, Hough-Neidig A, Chen Y, et al. GpsB Coordinates Cell Division and Cell Surface Decoration by Wall Teichoic Acids in *Staphylococcus aureus*. Microbiol Spectr. 2022;10(3):e0141322.

7. Sacco MD, Hammond LR, Noor RE, Bhattacharya D, McKnight L, Madsen JJ, et al. *Staphylococcus aureus* FtsZ and PBP4 bind to the conformationally dynamic N-terminal domain of GpsB. eLife. 2024.

8. Hammond LR, White ML, Eswara PJ. ¡vIVA la DivIVA! J Bacteriol. 2019;201(21).

9. Halbedel S, Lewis RJ. Structural basis for interaction of DivIVA/GpsB proteins with their ligands. Mol Microbiol. 2019;111(6):1404–15.

10. Barbuti MD, Myrbraten IS, Morales Angeles D, Kjos M. The cell cycle of Staphylococcus aureus: An updated review. Microbiologyopen. 2023;12(1):e1338.

11. Ramos-Leon F, Ramamurthi KS. Staphylococcus aureus as an emerging model to study bacterial cell division. J Biol Chem. 2025;301(7):110343.

12. Bartlett TM, Sisley TA, Mychack A, Walker S, Baker RW, Rudner DZ, et al. FacZ is a GpsB-interacting protein that prevents aberrant division-site placement in Staphylococcus aureus. Nat Microbiol. 2024;9(3):801–13.

13. Bhattacharya D, King A, McKnight L, Horigian P, Eswara PJ. GpsB interacts with FtsZ in multiple species and may serve as an accessory Z-ring anchor. Mol Biol Cell. 2025:mbcE24070302.

14. Sutton JAF, Cooke M, Tinajero-Trejo M, Wacnik K, Salamaga B, Portman-Ross C, et al. The roles of GpsB and DivIVA in Staphylococcus aureus growth and division. Front Microbiol. 2023;14:1241249.

15. Costa SF, Saraiva BM, Veiga H, Marques LB, Schaper S, Sporniak M, et al. The role of GpsB in Staphylococcus aureus cell morphogenesis. mBio. 2024;15(3):e0323523.

16. Fleurie A, Manuse S, Zhao C, Campo N, Cluzel C, Lavergne JP, et al. Interplay of the serine/threonine-kinase StkP and the paralogs DivIVA and GpsB in pneumococcal cell elongation and division. PLoS Genet. 2014;10(4):e1004275.

17. Pompeo F, Foulquier E, Serrano B, Grangeasse C, Galinier A. Phosphorylation of the cell division protein GpsB regulates PrkC kinase activity through a negative feedback loop in Bacillus subtilis. Mol Microbiol. 2015;97(1):139–50.

18. Kelliher JL, Grunenwald CM, Abrahams RR, Daanen ME, Lew CI, Rose WE, et al. PASTA kinase-dependent control of peptidoglycan synthesis via ReoM is required for cell wall stress responses, cytosolic survival, and virulence in Listeria monocytogenes. PLoS Pathog. 2021;17(10):e1009881.

19. VanZeeland NE, Schultz KM, Klug CS, Kristich CJ. Multisite Phosphorylation Regulates GpsB Function in Cephalosporin Resistance of Enterococcus faecalis. J Mol Biol. 2023;435(18):168216.

20. Rothe P, Wamp S, Rosemeyer L, Rismondo J, Doellinger J, Grundling A, et al. Cytosolic Factors Controlling PASTA Kinase-Dependent ReoM Phosphorylation. Mol Microbiol. 2024;122(4):514–33.

21. Stauberova V, Kubesa B, Joseph M, Benedet M, Furlan B, Buriankova K, et al. GpsB Coordinates StkP Signaling as a PASTA Kinase Adaptor in Streptococcus pneumoniae Cell Division. J Mol Biol. 2024;436(22):168797.

22. Junker S, Maabeta S, Otto A, Michalik S, Morgenroth F, Gerth U, et al. Spectral Library Based Analysis of Arginine Phosphorylations in Staphylococcus aureus. Mol Cell Proteomics. 2018;17(2):335–48.

23. Debarbouille M, Dramsi S, Dussurget O, Nahori MA, Vaganay E, Jouvion G, et al. Characterization of a serine/threonine kinase involved in virulence of Staphylococcus aureus. J Bacteriol. 2009;191(13):4070–81.

24. Beltramini AM, Mukhopadhyay CD, Pancholi V. Modulation of cell wall structure and antimicrobial susceptibility by a Staphylococcus aureus eukaryote-like serine/threonine kinase and phosphatase. Infect Immun. 2009;77(4):1406–16.

25. Ohlsen K, Donat S. The impact of serine/threonine phosphorylation in Staphylococcus aureus. Int J Med Microbiol. 2010;300(2-3):137–41.

26. Jarick M, Bertsche U, Stahl M, Schultz D, Methling K, Lalk M, et al. The serine/threonine kinase Stk and the phosphatase Stp regulate cell wall synthesis in Staphylococcus aureus. Sci Rep. 2018;8(1):13693.

27. Huemer M, Mairpady Shambat S, Hertegonne S, Bergada-Pijuan J, Chang CC, Pereira S, et al. Serine-threonine phosphoregulation by PknB and Stp contributes to quiescence and antibiotic tolerance in Staphylococcus aureus. Sci Signal. 2023;16(766):eabj8194.

28. Dunsmore CJ, Miller K, Blake KL, Patching SG, Henderson PJ, Garnett JA, et al. 2-Aminotetralones: novel inhibitors of MurA and MurZ. Bioorg Med Chem Lett. 2008;18(5):1730–4.

29. Blake KL, O’Neill AJ, Mengin-Lecreulx D, Henderson PJ, Bostock JM, Dunsmore CJ, et al. The nature of Staphylococcus aureus MurA and MurZ and approaches for detection of peptidoglycan biosynthesis inhibitors. Mol Microbiol. 2009;72(2):335–43.

30. Wamp S, Rutter ZJ, Rismondo J, Jennings CE, Moller L, Lewis RJ, et al. PrkA controls peptidoglycan biosynthesis through the essential phosphorylation of ReoM. Elife. 2020;9.

31. Sun Y, Hurlimann S, Garner E. Growth rate is modulated by monitoring cell wall precursors in Bacillus subtilis. Nat Microbiol. 2023;8(3):469–80.

32. Hall CL, Tschannen M, Worthey EA, Kristich CJ. IreB, a Ser/Thr kinase substrate, influences antimicrobial resistance in Enterococcus faecalis. Antimicrob Agents Chemother. 2013;57(12):6179–86.

33. Minton NE, Djoric D, Little J, Kristich CJ. GpsB Promotes PASTA Kinase Signaling and Cephalosporin Resistance in Enterococcus faecalis. J Bacteriol. 2022;204(10):e0030422.

34. Tsui HT, Joseph M, Zheng JJ, Perez AJ, Manzoor I, Rued BE, et al. Negative regulation of MurZ and MurA underlies the essentiality of GpsB- and StkP-mediated protein phosphorylation in Streptococcus pneumoniae D39. Mol Microbiol. 2023;120(3):351–83.

35. Canova MJ, Baronian G, Brelle S, Cohen-Gonsaud M, Bischoff M, Molle V. A novel mode of regulation of the Staphylococcus aureus Vancomycin-resistance-associated response regulator VraR mediated by Stk1 protein phosphorylation. Biochem Biophys Res Commun. 2014;447(1):165–71.

36. Kuroda M, Kuroda H, Oshima T, Takeuchi F, Mori H, Hiramatsu K. Two-component system VraSR positively modulates the regulation of cell-wall biosynthesis pathway in Staphylococcus aureus. Mol Microbiol. 2003;49(3):807–21.

37. Cleverley RM, Rismondo J, Lockhart-Cairns MP, Van Bentum PT, Egan AJ, Vollmer W, et al. Subunit Arrangement in GpsB, a Regulator of Cell Wall Biosynthesis. Microb Drug Resist. 2016;22(6):446–60.

38. Cleverley RM, Rutter ZJ, Rismondo J, Corona F, Tsui HT, Alatawi FA, et al. The cell cycle regulator GpsB functions as cytosolic adaptor for multiple cell wall enzymes. Nat Commun. 2019;10(1):261.

39. Baral R, Ho K, Kumar RP, Hopkins JB, Watkins MB, LaRussa S, et al. A general mechanism for initiating the bacterial general stress response. Elife. 2025;13.

40. Miksys A, Fu L, Madej MG, Guerreiro DN, Kaltwasser S, Conway M, et al. Molecular insights into intra-complex signal transmission during stressosome activation. Commun Biol. 2022;5(1):621.

41. Khadka R, Maravich B, Demarest N, Hartwig M, Tom A, Das NK, et al. Stressosome-independent but RsbT-dependent environmental stress sensing in Bacillus subtilis. Nat Commun. 2025;16(1):1591.

42. Mascari CA, Little JL, Kristich CJ. PASTA-kinase-mediated signaling drives accumulation of the peptidoglycan synthesis protein MurAA to promote cephalosporin resistance in Enterococcus faecalis. Mol Microbiol. 2023;120(6):811–29.

43. Battesti A, Bouveret E. The bacterial two-hybrid system based on adenylate cyclase reconstitution in Escherichia coli. Methods. 2012;58(4):325–34.

44. Adedeji-Olulana AF, Wacnik K, Lafage L, Pasquina-Lemonche L, Tinajero-Trejo M, Sutton JAF, et al. Two codependent routes lead to high-level MRSA. Science. 2024;386(6721):573–80.

45. Nagarajan SN, Lenoir C, Grangeasse C. Recent advances in bacterial signaling by serine/threonine protein kinases. Trends Microbiol. 2022;30(6):553–66.

46. Mascari CA, Djoric D, Kristich CJ. PASTA kinase signaling regulates peptidoglycan synthesis in Enterococcus faecalis by direct inhibition of UDP-N-acetylglucosamine 1-carboxyvinyl transferase activity. mBio. 2025;16(5):e0059325.

47. O’Reilly FJ, Graziadei A, Forbrig C, Bremenkamp R, Charles K, Lenz S, et al. Protein complexes in cells by AI-assisted structural proteomics. Mol Syst Biol. 2023;19(4):e11544.

48. Winkler ME, Joseph M, Tsui HT. Secrets of getting started: Regulation of the first committed step of peptidoglycan synthesis by protein phosphorylation in Enterococcus and other Gram-positive bacteria. Mol Microbiol. 2023;120(6):805–10.

49. Petek M, Baebler S, Kuzman D, Rotter A, Podlesek Z, Gruden K, et al. Revealing fosfomycin primary effect on Staphylococcus aureus transcriptome: modulation of cell envelope biosynthesis and phosphoenolpyruvate induced starvation. BMC Microbiol. 2010;10:159.

50. Weiss A, Jackson JK, Shaw LN, Skaar EP. Screening transcriptional connections in Staphylococcus aureus using high-throughput transduction of bioluminescent reporter plasmids. Microbiology (Reading). 2022;168(4).

51. Utaida S, Dunman PM, Macapagal D, Murphy E, Projan SJ, Singh VK, et al. Genome-wide transcriptional profiling of the response of Staphylococcus aureus to cell-wall-active antibiotics reveals a cell-wall-stress stimulon. Microbiology (Reading). 2003;149(Pt 10):2719–32.

52. Rismondo J, Bender JK, Halbedel S. Suppressor Mutations Linking gpsB with the First Committed Step of Peptidoglycan Biosynthesis in Listeria monocytogenes. J Bacteriol. 2017;199(1).

53. Hussein M, L. Neville S, Zhao J, R. Monk I, Wilksch J, Crawford S, et al. Metabolic reprogramming enables transient survival and envelope remodelling in Staphylococcus aureus under dalbavancin treatment. npj Antimicrobials and Resistance. 2026.

54. Sisley TA, Park Y, Santiago AG, Beroual W, Sobolewski IA, Lee W, et al. AuxB interacts directly with GpsB and PknB to coordinate cell envelope processes that contribute to intrinsic antibiotic resistance in Staphylococcus aureus. mBio. 2025;16(10):e0185825.

55. Kelliher JL, Daanen ME, Sauer J-D. GpsB control of PASTA kinase activity in <em>Listeria monocytogenes</em> influences peptidoglycan synthesis during cell wall stress and cytosolic survival. bioRxiv. 2023:2023.06.12.544644.

56. Graham JW, Lei MG, Lee CY. Trapping and identification of cellular substrates of the Staphylococcus aureus ClpC chaperone. J Bacteriol. 2013;195(19):4506–16.

57. Fernandes PB, Reed P, Monteiro JM, Pinho MG. Revisiting the Role of VraTSR in Staphylococcus aureus Response to Cell Wall-Targeting Antibiotics. J Bacteriol. 2022;204(8):e0016222.

58. Fey PD, Endres JL, Yajjala VK, Widhelm TJ, Boissy RJ, Bose JL, et al. A genetic resource for rapid and comprehensive phenotype screening of nonessential Staphylococcus aureus genes. mBio. 2013;4(1):e00537–12.

59. Kreiswirth BN, Lofdahl S, Betley MJ, O’Reilly M, Schlievert PM, Bergdoll MS, et al. The toxic shock syndrome exotoxin structural gene is not detectably transmitted by a prophage. Nature. 1983;305(5936):709–12.

60. Luong TT, Lee CY. The arl locus positively regulates Staphylococcus aureus type 5 capsule via an mgrA-dependent pathway. Microbiology (Reading). 2006;152(Pt 10):3123–31.

61. Waldo GS, Standish BM, Berendzen J, Terwilliger TC. Rapid protein-folding assay using green fluorescent protein. Nat Biotechnol. 1999;17(7):691–5.

62. Chen X, Zaro JL, Shen WC. Fusion protein linkers: property, design and functionality. Adv Drug Deliv Rev. 2013;65(10):1357–69.

63. Madeira F, Madhusoodanan N, Lee J, Eusebi A, Niewielska A, Tivey ARN, et al. The EMBL-EBI Job Dispatcher sequence analysis tools framework in 2024. Nucleic Acids Res. 2024;52(W1):W521–W5.

64. Waterhouse AM, Procter JB, Martin DM, Clamp M, Barton GJ. Jalview Version 2--a multiple sequence alignment editor and analysis workbench. Bioinformatics. 2009;25(9):1189–91.

