## Supplementary material for "*Staphylococcus aureus* GpsB, Stk, and VraSR form a cell wall surveillance network to maintain cell envelope integrity and mediate antibiotic resistance": Tables S1-S2 and Figures S1-S4

### I. Supplemental methods

#### Plasmid and Strain Construction

### II. Table and Figure Legends

**Table S1:** Strains used in this study

**Table S2:** Oligonucleotides used in this study

**Fig. S1: GpsB stability is not affected by phosphosite mutations.** GpsB protein abundance in strains from Fig. 1B-D was investigated via western blot using anti-GpsB antibody. SigA level, probed using anti-SigA antibody, served as loading control. Representative blots of biological triplicates are shown.

**Fig. S2: GpsB phosphomodulation is important for fosfomycin resistance.** Fosfomycin (Fos) influence on the growth of *S. aureus* JE2 and Δ*gpsB* strains harboring either empty vector (EV) or *gpsB*/phosphomutants in liquid culture (**A**) or solid medium (**B**). Strains used: JE2 + EV (WT, LM253), Δ*gpsB* + EV (LM254), Δ*gpsB* + *gpsB* (LM255), Δ*gpsB* + AAA (LM319), and Δ*gpsB* + EEE (LM320). Growth without or with Fos at concentrations indicated was monitored for 10 h (A) or after overnight incubation (B) at 37 °C. Representative growth curves and plate pictures are shown, n=3. (C) Summary of results provided in panels A and B. In the absence of either GpsB or Stk/Stp regulation (*of GpsB), cells are unable to achieve maximal Fos resistance.

**Fig. S3: *murZ* promoter activity is diminished in the absence of GpsB.** (**A**) Bacterial two-hybrid assay results. Interactions between T18 fusions of MurA, MurZ and ReoM were tested against T25 fusions of MurA, MurZ, ReoM and ReoY. Representative plate images of three independent biological replicates are shown. (**B-C**) Growth ($OD_{600}$; top panels) and luminescence (arbitrary units; bottom panels) readings of strains indicated below, harboring $P_{murZ}$-*lux* transcriptional reporter, were monitored for 10 h at 37°C in the absence or presence of fosfomycin (2.5 μg/ml). Reporter strains used: WT JE2 (LM329), Δ*gpsB* (LM330), Δ*stk* (LM331), Δ*stk* Δ*gpsB* (LM332), Δ*murA* (LM333), and Δ*murA* Δ*gpsB* (LM334). Representative graphs are shown, n=3.

**Fig. S4: GpsB and VraSR coordinate to confer fosfomycin resistance.** (**A-B**) Growth curves ($OD_{600}$) of WT *S. aureus* (JE2), Δ*gpsB* (LM252), Δ*vraS* (LM336), Δ*vraS* Δ*gpsB* (LM337), Δ*vraR* (LM338), and Δ*vraR* Δ*gpsB* (LM339) strains were grown in the absence or presence of fosfomycin at concentrations indicated for 10 h at 37 °C. (**C-D**) Growth curves of WT *S. aureus* JE2, *gpsB* single-deletion, and *gpsB vraR* or *gpsB vraS* double-deletion strains harboring either empty vector (EV) or *gpsB*/phosphomutants were grown in the absence or presence of fosfomycin at indicated concentrations for 10 h at 37 °C. Strains used: JE2 + EV (WT, LM253), Δ*gpsB* + EV (LM254), Δ*gpsB* + *gpsB* (LM255), Δ*vraR* Δ*gpsB* + EV (LM344), Δ*vraR* Δ*gpsB* + *gpsB* (LM345), Δ*vraR* Δ*gpsB* + T75E (LM346), Δ*vraR* Δ*gpsB* + T75A (LM347), Δ*vraS* Δ*gpsB* + EV (LM340), Δ*vraS* Δ*gpsB* + *gpsB* (LM341), Δ*vraS* Δ*gpsB* + T75E (LM342), and Δ*vraS* Δ*gpsB* + T75A (LM343). Representative graphs are shown, n=3.

### I. Supplemental methods

#### Plasmid and Strain Construction

Standard molecular biology protocols were employed to construct plasmids and *Escherichia coli* DH5α was used as primary plasmid host. The insert sequences in all plasmids generated were confirmed via either Sanger or whole plasmid sequencing (GeneWiz/Azenta).

<u>*gpsB/phosphomutant* complementation:</u>

The *gpsB* coding region was PCR amplified from JE2 genomic DNA using primers oLM48 and oLM49, which introduce homology to the pCL15 multiple cloning site along with the ribosome binding site and start codon (oLM48) or stop codon (oLM49). This fragment was then integrated by Gibson assembly into pCL15 linearized with primers oLM50/oLM51 and subsequently treated with DpnI enzyme, yielding the plasmid pPE45. The empty vector (pCL15) was electroporated into *S. aureus* RN4220 Δ*gpsB* to generate LM74 and LM75 strains respectively. Similarly, pCL15 or pPE45 was introduced into *S. aureus* JE2 *gpsB*::*kan* via phage transduction (bacteriophage Φ11 with selection on 10 µg/mL chloramphenicol) to generate strains LM254 and LM255 respectively. QuikChange (Agilent) site-directed mutagenesis was used to introduce desired phosphomutation(s) in pPE45 using the primer pairs listed in Table S2. Plasmids from *E. coli* DH5a strains LM211-LM230 were purified and electroporated into *S. aureus* RN4220 Δ*gpsB* resulting in strains LM235-LM251. RN4220 strains harboring desired plasmids were then introduced into JE2 or different mutant backgrounds using bacteriophage Φ11 and selection on 10 µg/mL chloramphenicol. All the strains generated for this study are listed in Table S1.

<u>*murZ* promoter reporter:</u>

For $P_{murZ}$-*lux* transcriptional reporter, the *murZ* promoter region was amplified from JE2 genomic DNA with primers oLM171/oLM172 and assembled by Gibson assembly into pXEN-1, yielding pLM124. This reporter plasmid was electroporated into RN4220 (LM328) and subsequently transduced into the WT, Δ*gpsB*, Δ*stk,* Δ*stk* Δ*gpsB*, Δ*murA*, and Δ*murA* Δ*gpsB* strains of JE2 background (LM329–LM334).

<u>Bacterial two-hybrid plasmids:</u>

For bacterial two-hybrid analysis, *gpsB* (WT and T75E/T75A point mutants) was PCR-amplified with primers appending a C-terminal flexible (GSAGSAAGSGEF (1, 2)) linker (oLM60/oLM61 for pKNT25; oLM80/oLM81 for pUT18) and cloned by Gibson assembly into pKNT25 and pUT18 linearized with oLM62/oLM63 and oLM78/oLM79, respectively, yielding pLM94/pLM95 (WT) and the corresponding pLM108/pLM111 (T75E) and pLM112/pLM113 (T75A) plasmids. *stk* and *stp* were amplified with primers oLM97/oLM100 (EcoRI/XhoI) and cloned into pEB354/pEB355, yielding pLM107 and pLM116 respectively. *murA* (oLM163/164), *murZ* (oLM165/166), *reoM* (oLM167/168), and *reoY* (oLM169/170) were amplified from JE genomic DNA and cloned using restriction digest (EcoRI/XhoI) method (or Gibson assembly with homology to pEB354 for *murZ*) and cloned into pEB354, yielding pLM120-pLM123 plasmids. *vraS* (oLM152/oLM153) and *vraR* (oLM189/190) were amplified from JE2 genomic DNA and cloned using restriction enzymes into pEB354 (EcoRI/XhoI) and pKNT25 (PstI/BamHI). *pbp1*, *pbp2*, and *pbp3* were similarly amplified from JE2 genomic DNA and cloned into pEB354: *pbp1* with primers BTH1/BTH2 (EcoRI/XhoI), *pbp2* with primers BTH3/BTH4 (PstI/XhoI), and *pbp3* with primers BTH5/BTH6 (EcoRI/XhoI), yielding pLH34, pLH37, and pLH35, respectively. *mecA*, which encodes PBP2a, was amplified from JE2 chromosomal DNA with Gibson primers oLM141/oLM142 and cloned into pEB354, yielding pLM125 (strain LM335). pEB354-based *tarG* BTH plasmid (pSKB1) was generated in our laboratory for a previous study (3).

Protein purification:

We used the pET28a-based construct of his6-*gpsB* (pLM117; (4)) as site-directed mutagenesis template to introduce T75E substitution (pLM118). Both pLM117 and pLM118 were transformed into *E. coli* BL21 (DE3) strain for protein purification.

**Table S1:** Strains used in this study

| Species | Strain | Genotype | Plasmid | Source |
|---|---|---|---|---|
| *S. aureus* strains | | | | |
| *S. aureus* | RN4220 | Wild type | | (5) |
| *S. aureus* | JE2 | Wild type | | (6) |
| *S. aureus* | SJF4925 | SH1000 Δ*gpsB::kan* | | (7) |
| *S. aureus* | LM74 | RN4220 Δ*gpsB* pCL15 | pCL15 | This study; (8) |
| *S. aureus* | LM75 | RN4220 Δ*gpsB* pCL15-*gpsB* | pPE45 | This study; (4) |
| *S. aureus* | LM235 | RN4220 Δ*gpsB* pCL15-*gpsB-T21E* | pLM66 | This study |
| *S. aureus* | LM236 | RN4220 Δ*gpsB* pCL15-*gpsB-T75E* | pLM67 | This study |
| *S. aureus* | LM237 | RN4220 Δ*gpsB* pCL15-*gpsB-S76E* | pLM68 | This study |
| *S. aureus* | LM238 | RN4220 Δ*gpsB* pCL15-*gpsB-S85E* | pLM69 | This study |
| *S. aureus* | LM239 | RN4220 Δ*gpsB* pCL15-*gpsB-T93E* | pLM70 | This study |
| *S. aureus* | LM240 | RN4220 Δ*gpsB* pCL15-*gpsB-S105E* | pLM71 | This study |
| *S. aureus* | LM241 | RN4220 Δ*gpsB* pCL15-*gpsB-T21A* | pLM72 | This study |
| *S. aureus* | LM242 | RN4220 Δ*gpsB* pCL15-*gpsB-T75A* | pLM73 | This study |
| *S. aureus* | LM243 | RN4220 Δ*gpsB* pCL15-*gpsB-S76A* | pLM74 | This study |
| *S. aureus* | LM244 | RN4220 Δ*gpsB* pCL15-*gpsB-EEE* (T21E S76E T75E) | pLM78 | This study |
| *S. aureus* | LM245 | RN4220 Δ*gpsB* pCL15-*gpsB-EAA* (T21E S76A T75A) | pLM79 | This study |
| *S. aureus* | LM246 | RN4220 Δ*gpsB* pCL15-*gpsB-AEE* (T21A S76E T75E) | pLM80 | This study |
| *S. aureus* | LM247 | RN4220 Δ*gpsB* pCL15-*gpsB-AAA* (T21A S76A T75A) | pLM81 | This study |
| *S. aureus* | LM248 | RN4220 Δ*gpsB* pCL15-*gpsB-AE* (T75A S76E) | pLM82 | This study |
| *S. aureus* | LM249 | RN4220 Δ*gpsB* pCL15-*gpsB-EA* (T75E S76A) | pLM83 | This study |
| *S. aureus* | LM250 | RN4220 Δ*gpsB* pCL15-*gpsB-EE* (T75E S76E) | pLM84 | This study |
| *S. aureus* | LM251 | RN4220 Δ*gpsB* pCL15-*gpsB-AA* (T75A S76A) | pLM85 | This study |
| *S. aureus* | LM252 | JE2 Δ*gpsB::kan* | | This study; (7) |
| *S. aureus* | LM253 | JE2 pCL15 | pCL15 | This study |
| *S. aureus* | LM254 | JE2 Δ*gpsB::kan* pCL15 | pCL15 | This study |
| *S. aureus* | LM255 | JE2 Δ*gpsB::kan* pCL15-*gpsB* | pPE45 | This study |
| *S. aureus* | LM256 | JE2 Δ*gpsB::kan* pCL15-*gpsB-T75E* | pLM67 | This study |
| *S. aureus* | LM257 | JE2 Δ*gpsB::kan* pCL15-*gpsB-T75A* | pLM73 | This study |

| | | | | |
|---|---|---|---|---|
| *S. aureus* | LM260 | JE2 *stk::erm* | | This study (NTML) |
| *S. aureus* | LM310 | JE2 Δ*gpsB::kan stk::erm* | | This study |
| *S. aureus* | LM311 | JE2 *reoY::erm* | | This study (NTML) |
| *S. aureus* | LM312 | JE2 *reoY::erm gpsB::kan* | | This study |
| *S. aureus* | LM313 | JE2 *stp::erm* | | This study (NTML) |
| *S. aureus* | LM314 | JE2 *stp::erm gpsB::kan* | | This study |
| *S. aureus* | LM315 | JE2 *stk::erm gpsB::kan* pCL15 | pCL15 (EV) | This study |
| *S. aureus* | LM316 | JE2 *stk::erm gpsB::kan* pCL15-*gpsB* | pPE45 | This study |
| *S. aureus* | LM317 | JE2 *stp::erm gpsB::kan* pCL15 | pCL15 (EV) | This study |
| *S. aureus* | LM318 | JE2 *stp::erm gpsB::kan* pCL15-*gpsB* | pPE45 | This study |
| *S. aureus* | LM319 | JE2 *gpsB::kan* pCL15-*gpsB-AAA* | pLM81 | This study |
| *S. aureus* | LM320 | JE2 *gpsB::kan* pCL15-*gpsB-EEE* | pLM78 | This study |
| *S. aureus* | LM325 | JE2 *murA::erm* | | This study (NTML) |
| *S. aureus* | LM326 | JE2 *murA::erm gpsB::kan* | | This study |
| *S. aureus* | LM328 | RN4220 $P_{murZ}$-*lux* reporter | pLM124 | This study |
| *S. aureus* | LM329 | JE2 $P_{murZ}$-*lux* reporter | pLM124 | This study |
| *S. aureus* | LM330 | JE2 *gpsB::kan* $P_{murZ}$-*lux* reporter | pLM124 | This study |
| *S. aureus* | LM331 | JE2 *stk::erm* $P_{murZ}$-*lux* reporter | pLM124 | This study |
| *S. aureus* | LM332 | JE2 *stk::erm gpsB::kan* $P_{murZ}$-*lux* reporter | pLM124 | This study |
| *S. aureus* | LM333 | JE2 *murA::erm* $P_{murZ}$-*lux* reporter | pLM124 | This study |
| *S. aureus* | LM334 | JE2 *murA::erm gpsB::kan* $P_{murZ}$-*lux* reporter | pLM124 | This study |
| *S. aureus* | LM336 | JE2 *vraS::erm* | | This study (NTML) |
| *S. aureus* | LM337 | JE2 *vraS::erm gpsB::kan* | | This study |
| *S. aureus* | LM338 | JE2 *vraR::erm* | | This study (NTML) |
| *S. aureus* | LM339 | JE2 *vraR::erm gpsB::kan* | | This study |
| *S. aureus* | LM340 | JE2 *vraS::erm gpsB::kan* pCL15 | pCL15 (EV) | This study |
| *S. aureus* | LM341 | JE2 *vraS::erm gpsB::kan* pCL15-*gpsB* | pPE45 | This study |
| *S. aureus* | LM342 | JE2 *vraS::erm gpsB::kan* pCL15-*gpsB-T75E* | pLM67 | This study |
| *S. aureus* | LM343 | JE2 *vraS::erm gpsB::kan* pCL15-*gpsB-T75A* | pLM73 | This study |
| *S. aureus* | LM344 | JE2 *vraR::erm gpsB::kan* pCL15 | pCL15 (EV) | This study |
| *S. aureus* | LM345 | JE2 *vraR::erm gpsB::kan* pCL15-*gpsB* | pPE45 | This study |
| *S. aureus* | LM346 | JE2 *vraR::erm gpsB::kan* pCL15-*gpsB-T75E* | pLM67 | This study |
| *S. aureus* | LM347 | JE2 *vraR::erm gpsB::kan* pCL15-*gpsB-T75A* | pLM73 | This study |
| *E. coli* strains | | | | |
| *E. coli* | LM211 | DH5α harboring pCL15-*gpsB-T21E* | pLM66 | This study |
| *E. coli* | LM212 | DH5α harboring pCL15-*gpsB-T75E* | pLM67 | This study |
| *E. coli* | LM213 | DH5α harboring pCL15-*gpsB-S76E* | pLM68 | This study |
| *E. coli* | LM214 | DH5α harboring pCL15-*gpsB-S85E* | pLM69 | This study |
| *E. coli* | LM215 | DH5α harboring pCL15-*gpsB-T93E* | pLM70 | This study |
| *E. coli* | LM216 | DH5α harboring pCL15-*gpsB-S105E* | pLM71 | This study |
| *E. coli* | LM217 | DH5α harboring pCL15-*gpsB-T21A* | pLM72 | This study |
| *E. coli* | LM218 | DH5α harboring pCL15-*gpsB-T75A* | pLM73 | This study |
| *E. coli* | LM219 | DH5α harboring pCL15-*gpsB-S76A* | pLM74 | This study |
| *E. coli* | LM220 | DH5α harboring pCL15-*gpsB-S85A* | pLM75 | This study |
| *E. coli* | LM221 | DH5α harboring pCL15-*gpsB-T93A* | pLM76 | This study |

| | | | | |
|---|---|---|---|---|
| *E. coli* | LM222 | DH5α harboring pCL15-*gpsB-S105A* | pLM77 | This study |
| *E. coli* | LM223 | DH5α harboring pCL15-*gpsB-EEE* | pLM78 | This study |
| *E. coli* | LM224 | DH5α harboring pCL15-*gpsB-EAA* | pLM79 | This study |
| *E. coli* | LM225 | DH5α harboring pCL15-*gpsB-AEE* | pLM80 | This study |
| *E. coli* | LM226 | DH5α harboring pCL15-*gpsB-AAA* | pLM81 | This study |
| *E. coli* | LM227 | DH5α harboring pCL15-*gpsB-AE* | pLM82 | This study |
| *E. coli* | LM228 | DH5α harboring pCL15-*gpsB-EA* | pLM83 | This study |
| *E. coli* | LM229 | DH5α harboring pCL15-*gpsB-EE* | pLM84 | This study |
| *E. coli* | LM230 | DH5α harboring pCL15-*gpsB-AA* | pLM85 | This study |
| *E. coli* | LM266 | DH5α harboring pKNT25-*gpsB-linker* (GpsB-T25) | pLM94 | This study |
| *E. coli* | LM267 | DH5α harboring pUT18-*gpsB-linker* (GpsB-T18) | pLM95 | This study |
| *E. coli* | LM279 | DH5α harboring pEB354-*stk* (T25-Stk) | pLM107 | This study |
| *E. coli* | LM280 | DH5α harboring pUT18-*gpsB-T75E-linker* (GpsB$^{T75E}$-T18) | pLM108 | This study |
| *E. coli* | LM283 | DH5α harboring pKNT25-*gpsB-T75E-linker* (GpsB$^{T75E}$-T25) | pLM111 | This study |
| *E. coli* | LM284 | DH5α harboring pKNT25-*gpsB-T75A-linker* (GpsB$^{T75A}$-T25) | pLM112 | This study |
| *E. coli* | LM285 | DH5α harboring pUT18-*gpsB-T75A-linker* (GpsB$^{T75A}$-T18) | pLM113 | This study |
| *E. coli* | LM288 | DH5α harboring pEB354-*stp* (T25-Stp) | pLM116 | This study |
| *E. coli* | LM289 | BL21(DE3) harboring pET28a-*gpsB* | pLM117 | (9) |
| *E. coli* | LM290 | BL21(DE3) harboring pET28a-*gpsB-T75E* | pLM118 | This study |
| *E. coli* | LM321 | DH5α harboring pEB354-*murA* (T25-MurA) | pLM120 | This study |
| *E. coli* | LM322 | DH5α harboring pEB354-*murZ* (T25-MurZ) | pLM121 | This study |
| *E. coli* | LM323 | DH5α harboring pEB354-*reoM* (T25-ReoM) | pLM122 | This study |
| *E. coli* | LM324 | DH5α harboring pEB354-*reoY* (T25-ReoY) | pLM123 | This study |
| *E. coli* | LM327 | DH5α harboring pXEN1-$P_{murZ}$-*lux* | pLM124 | This study |
| *E. coli* | LM348 | DH5α harboring pEB355-*murA* (T18-MurA) | pLM125 | This study |
| *E. coli* | LM349 | DH5α harboring pEB355-*murZ* (T18-MurZ) | pLM126 | This study |
| *E. coli* | LM350 | DH5α harboring pEB355-*reoM* (T18-ReoM) | pLM127 | This study |
| *E. coli* | LM351 | DH5α harboring pEB354-*vraR* (T25-VraR) | pLM128 | This study |
| *E. coli* | LM352 | DH5α harboring pKNT25-*vraS* (VraS-T25) | pLM129 | This study |
| *E. coli* | LH102 | DH5α harboring pEB354-*pbp1* (T25-PBP1) | pLH34 | This study |
| *E. coli* | LH103 | DH5α harboring pEB354-*pbpC* (T25-PBP3) | pLH35 | This study |
| *E. coli* | LH105 | DH5α harboring pEB354-*pbpB* (T25-PBP2) | pLH37 | This study |
| *E. coli* | LM335 | DH5α harboring pEB354-*mecA* (T25-PBP2a) | pLM125 | This study |
| *E. coli* | SKB1 | DH5α harboring pEB354-*tarG* (T25-TarG) | pSKB1 | (3) |

**Table S2:** Oligonucleotides used in this study

| | Primer # | Sequence (5' to 3') |
|---|---|---|
| T21A | oLH27 | atagccacgagccatcgctttttcaaaatctttttcataaatatcttttgc |
| | oLH28 | gcaaaagatatttatgaaaaagattttgaaaaagcgatggctcgtggctat |
| T21E | oLH29 | tctatagccacgagccatctctttttcaaaatctttttcataaatatcttttgctg |
| | oLH30 | cagcaaaagatatttatgaaaaagattttgaaaaagagatggctcgtggctataga |
| T75A | oLH31 | gtcctgaggtcttgatgctgctacacgtagtctta |
| | oLH32 | taagactacgtgtagcagcatcaagacctcaggac |
| T75E | oLH33 | cttttattgtcctgaggtcttgactctgctacacgtagtcttaattcttctaattc |
| | oLH34 | gaattagaagaattaagactacgtgtagcagagtcaagacctcaggacaataaaag |
| S76A | oLH35 | attgtcctgaggtcttgctgttgctacacgtagtc |
| | oLH36 | gactacgtgtagcaacagcaagacctcaggacaat |
| S76E | oLH37 | cttttattgtcctgaggtctctctgttgctacacgtagtcttaattcttctaattcttttt |
| | oLH38 | aaaaagaattagaagaattaagactacgtgtagcaacagagagacctcaggacaataaaag |
| S85E | oLH41 | tgtattattcgaagaaaaacttttattgtcctgaggtctctcctctgctacacgtagtcttaattcttctaattcttttttaagtttatg |
| | oLH42 | cataaacttaaaaaagaattagaagaattaagactacgtgtagcagaggagagacctcaggacaataaaagtttttcttcgaataataca |
| T93E | oLH49 | aatacgttttaaaatatctacattatttgaagactcatttgttgttgtattattcgaagaaaaacttttattgtcctgag |
| | oLH50 | ctcaggacaataaaagtttttcttcgaataatacaacaacaaatgagtcttcaaataatgtagatattttaaaacgtatt |
| S105E | oLH53 | tttaccaaatacagctttttctaagttctcaatacgttttaaaatatctacattatttgaagatgtatttgttgttgtatta |
| | oLH54 | taatacaacaacaaatacatcttcaaataatgtagatattttaaaacgtattgagaacttagaaaaagctgtatttggtaaa |
| T75A + | oLM1 | gactacgtgtagcagaggcaagacctcaggacaat |
| S76A | oLM2 | attgtcctgaggtcttgcctctgctacacgtagtc |
| T75E + | oLM3 | taagactacgtgtagcagcagagagacctcaggac |
| S76E | oLM4 | gtcctgaggtctctctgctgctacacgtagtctta |
| | oLM48 | atgtgtggaattgtgagcggataacaattaagcttACATAAGGAGGAACTACTATGTC |
| | oLM49 | cccggggatcctctagagtcgacctgcaggcatgcTTATTTACCAAATACAGCTTTTTCTAAG |
| | oLM50 | GCATGCCTGCAGGTCGACTC |
| | oLM51 | AAGCTTAATTGTTATCCGCTCACAATTC |
| | oLM52 | tatGGATCCgATGGATTGGATTTTACCAAT |
| | oLM53 | tgcgGAATTCgcTTTATCTACTCTAGAAGTATAGCTATG |
| | oLM60 | ccatgattacgccaagcttgatgtcagatgtttcattgaaattatc |
| | oLM61 | agagtcgacctgcaggcatgGAACTCGCCAGAACCAGCAGCGGAGCCAGCGGAACCTTTACCAAATACAGCTTTTTC |
| | oLM62 | catgcctgcaggtcgactc |
| | oLM63 | caagcttggcgtaatcatgg |
| | oLM78 | tgcaggtcgactctagagg |
| | oLM79 | ggcatgcaagcttggcgtaatc |
| | oLM80 | atgaccatgattacgccaagcttgcatgccatgtcagatgtttcattgaaattatc |
| | oLM81 | tacccggggatcctctagagtcgacctgcaGAACTCGCCAGAACCAGCAGCGGAGCCAGCGGAACCtttaccaaatacagctttttctaag |
| | oLM84 | ccccaggctttacacttt |
| | oLM85 | gcccactgcggaacg |
| | oLM86 | cggctgcggggatgc |
| | oLM97 | ATTGAATTCATGATAGGTAAAATAATAAATGAACG |
| | oLM98 | tgtgCTCGAGTTATACATCATCATAGCTGACTT |
| | oLM99 | tgtgGAATTCATGCTAGAGGCACAATTTT |
| | oLM100 | tgtgCTCGAGTCATACTTTATCACCTTCAATAG |
| | oLM141 | gcagggtcgactctagaattcgatatcaagatgaaaaagataaaaattgttccac |

| | |
|---|---|
| oLM142 | gaattgctcgaggtcgacggtatcgataagttattcatctatatcgtattttttattacc |
| oLM163 | tataGAATTCATGGATAAAATAGTAATCAAAGGTG |
| oLM164 | tataCTCGAGTTAATCGTTAATACGTTCAATGTCT |
| oLM165 | ctctagaattcgatatcaagatggctcaagaggtaataaaaataag |
| oLM166 | aggtcgacggtatcgataagctatacagtttccgtccaaatatc |
| oLM167 | tataGAATTCATGTCAGAGACACTCAATCA |
| oLM168 | tataCTCGAGTTAATTGTCATCTGCTAAGTCC |
| oLM169 | tataGAATTCATGGAAAACTTTGATAAAACAATGA |
| oLM170 | tataCTCGAGTTATTTATTTTGCTCTTTTAAATAGTAAGAAA |
| oLM171 | aaacgacggccagtgaattcaattgattgtgatataattataaggaaaatg |
| oLM172 | ttcatctgcaggatccccggatcatttatctcctttgtcc |
| oLM173 | ataaacgtgtgagtccagatttgtttat |
| oLM174 | ataaatgcagtaaaattaatcgagaggt |
| oLM175 | aagaagatttggatcctacatt |
| oLM176 | atttgtaacattattcaaatatggg |
| oLM177 | atttaataaaaggcttaaactgc |
| oLM178 | ttgatgaagaagatgactgt |
| oLM179 | tataGCATGCAAGCTTACATAAGGAGGAACTACTATGGATAAAATAGTAATCAAAGGTG |
| oLM180 | tataCTGCAGTTAATCGTTAATACGTTCAATGTCT |
| oLM189 | tataGAATTCATGACGATTAAAGTATTGTTTGT |
| oLM190 | tataCTCGAGCTATTGAATTAAATTATGTTGGAATG |
| BTH1 | AATAAGAATTCATGGCGAAGCAAAAAATTAAAATTAAAAAAAATAAAATAGG |
| BTH2 | AATAACTCGAGTTAGTCCGACTTATCCTTGTCAG |
| BTH3 | AATAACTGCAGATGACGGAAAACAAAGGATCTTCTCAGCC |
| BTH4 | AATAACTCGAGTTAGTTGAATATACCTGTTAATCC |
| BTH5 | AATAAGAATTCATGTTAAAAAGACTAAAAGAAAAATCAAATGATGAAATCG |
| BTH6 | AATAACTCGAG TTATTTGTCTTTGTCTTTATTTTTATCATCTTTACC |

### References

1. Waldo GS, Standish BM, Berendzen J, Terwilliger TC. Rapid protein-folding assay using green fluorescent protein. Nat Biotechnol. 1999;17(7):691-5.
2. Chen X, Zaro JL, Shen WC. Fusion protein linkers: property, design and functionality. Adv Drug Deliv Rev. 2013;65(10):1357-69.
3. Hammond LR, Sacco MD, Khan SJ, Spanoudis C, Hough-Neidig A, Chen Y, et al. GpsB Coordinates Cell Division and Cell Surface Decoration by Wall Teichoic Acids in *Staphylococcus aureus*. Microbiol Spectr. 2022;10(3):e0141322.
4. Eswara PJ, Brzozowski RS, Viola MG, Graham G, Spanoudis C, Trebino C, et al. An essential *Staphylococcus aureus* cell division protein directly regulates FtsZ dynamics. Elife. 2018;7.
5. Kreiswirth BN, Lofdahl S, Betley MJ, O'Reilly M, Schlievert PM, Bergdoll MS, et al. The toxic shock syndrome exotoxin structural gene is not detectably transmitted by a prophage. Nature. 1983;305(5936):709-12.
6. Fey PD, Endres JL, Yajjala VK, Widhelm TJ, Boissy RJ, Bose JL, et al. A genetic resource for rapid and comprehensive phenotype screening of nonessential Staphylococcus aureus genes. mBio. 2013;4(1):e00537-12.
7. Sutton JAF, Cooke M, Tinajero-Trejo M, Wacnik K, Salamaga B, Portman-Ross C, et al. The roles of GpsB and DivIVA in Staphylococcus aureus growth and division. Front Microbiol. 2023;14:1241249.
8. Luong TT, Lee CY. The arl locus positively regulates Staphylococcus aureus type 5 capsule via an mgrA-dependent pathway. Microbiology (Reading). 2006;152(Pt 10):3123-31.
9. Bhattacharya D, King A, McKnight L, Horigian P, Eswara PJ. GpsB interacts with FtsZ in multiple species and may serve as an accessory Z-ring anchor. Mol Biol Cell. 2025:mbcE24070302.

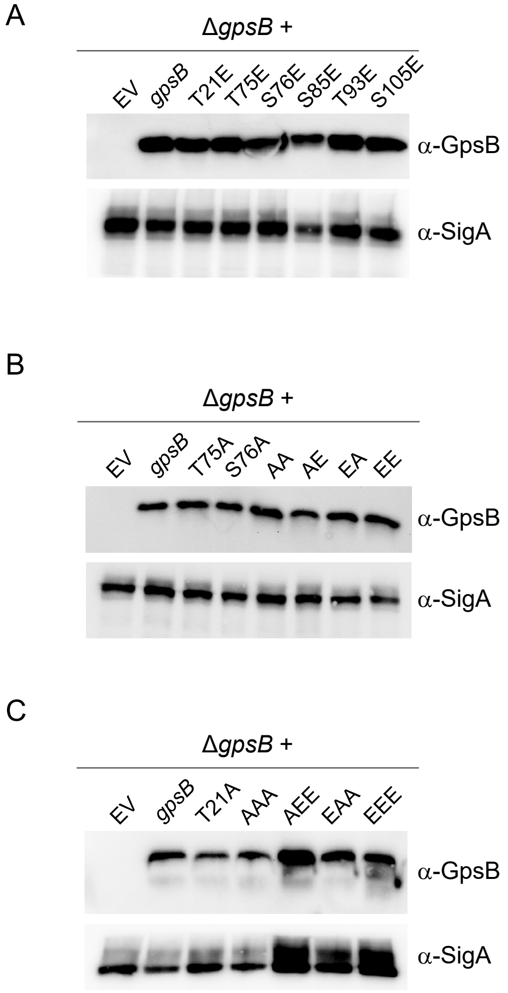
A
ΔgpsB +
EV
gpsB
T21E
T75E
S76E
S85E
T93E
S105E
α-GpsB
α-SigA
B
ΔgpsB +
EV
gpsB
T75A
S76A
AA
AE
EA
EE
α-GpsB
α-SigA
C
ΔgpsB +
EV
gpsB
T21A
AAA
AEE
EAA
EEE
α-GpsB
α-SigA

Figure S1

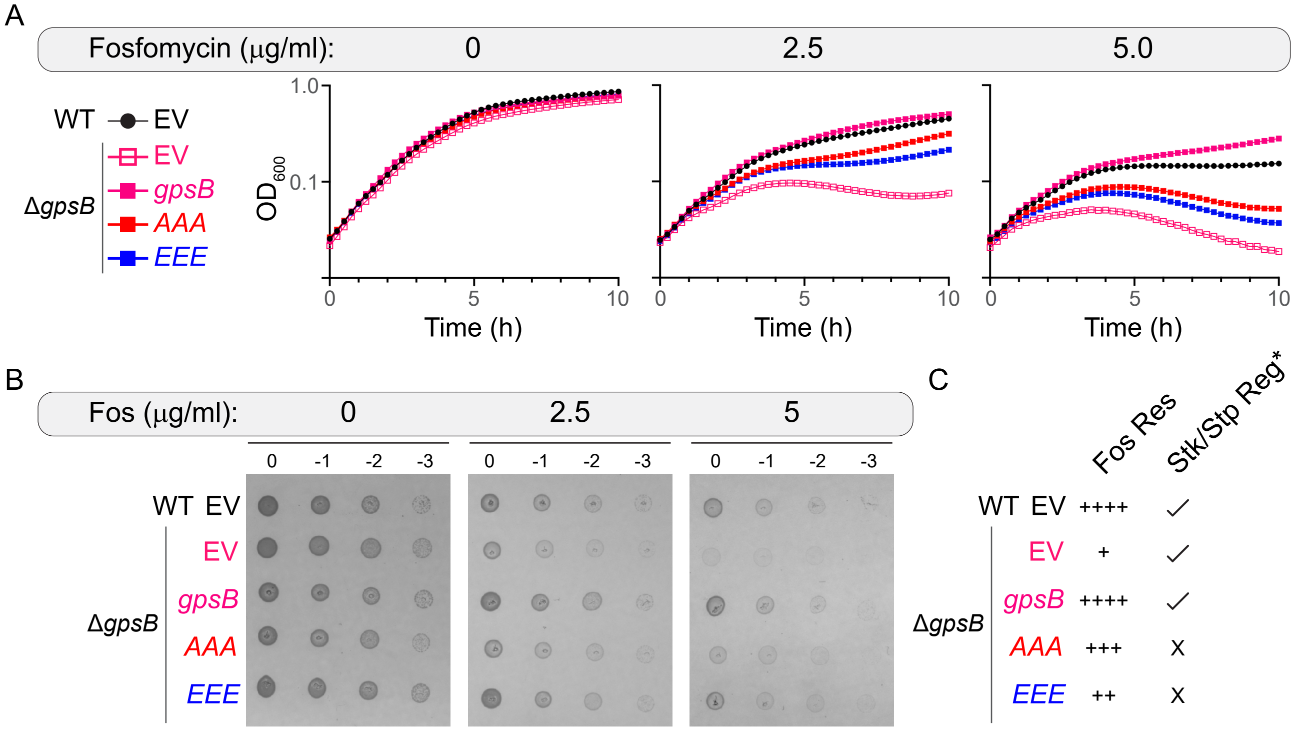
A
Fosfomycin (μg/ml): 0 2.5 5.0
WT EV
ΔgpsB EV gpsB AAA EEE
OD600
1.0
0.1
Time (h)
0 5 10
B
Fos (μg/ml): 0 2.5 5
0 -1 -2 -3
WT EV
ΔgpsB EV gpsB AAA EEE
C
Fos Res
Stk/Stp Reg*
WT EV ++++ ✓
ΔgpsB EV + ✓
gpsB ++++ ✓
AAA +++ X
EEE ++ X

Figure S2

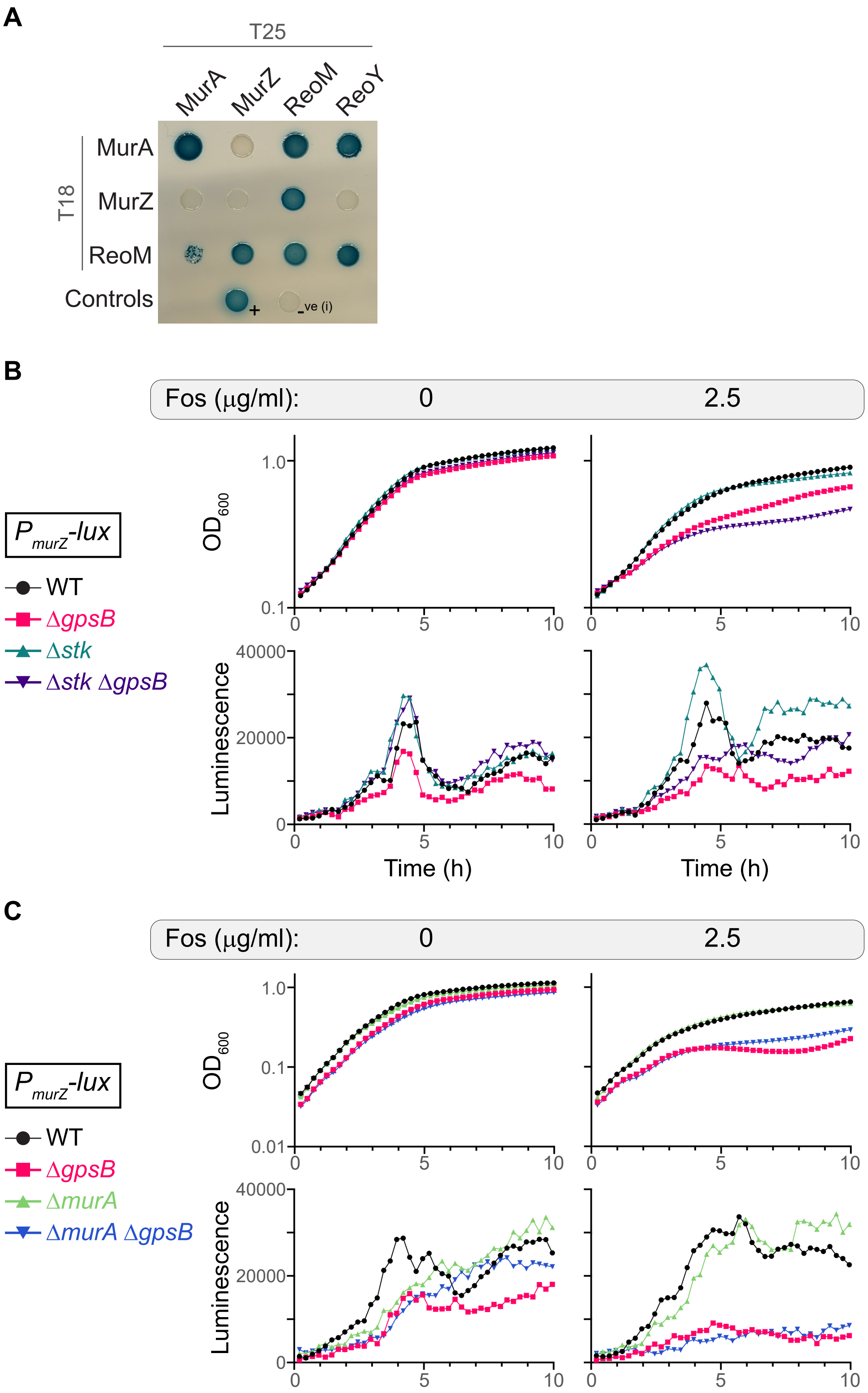
A
T25
MurA
MurZ
ReoM
ReoY
T18
MurA
MurZ
ReoM
Controls
+
-ve (i)
B
Fos (µg/ml):
0
2.5
$P_{murZ}$-lux
WT
ΔgpsB
Δstk
Δstk ΔgpsB
OD600
Luminescence
Time (h)
C
Fos (µg/ml):
0
2.5
$P_{murZ}$-lux
WT
ΔgpsB
ΔmurA
ΔmurA ΔgpsB
OD600
Luminescence

Figure S3

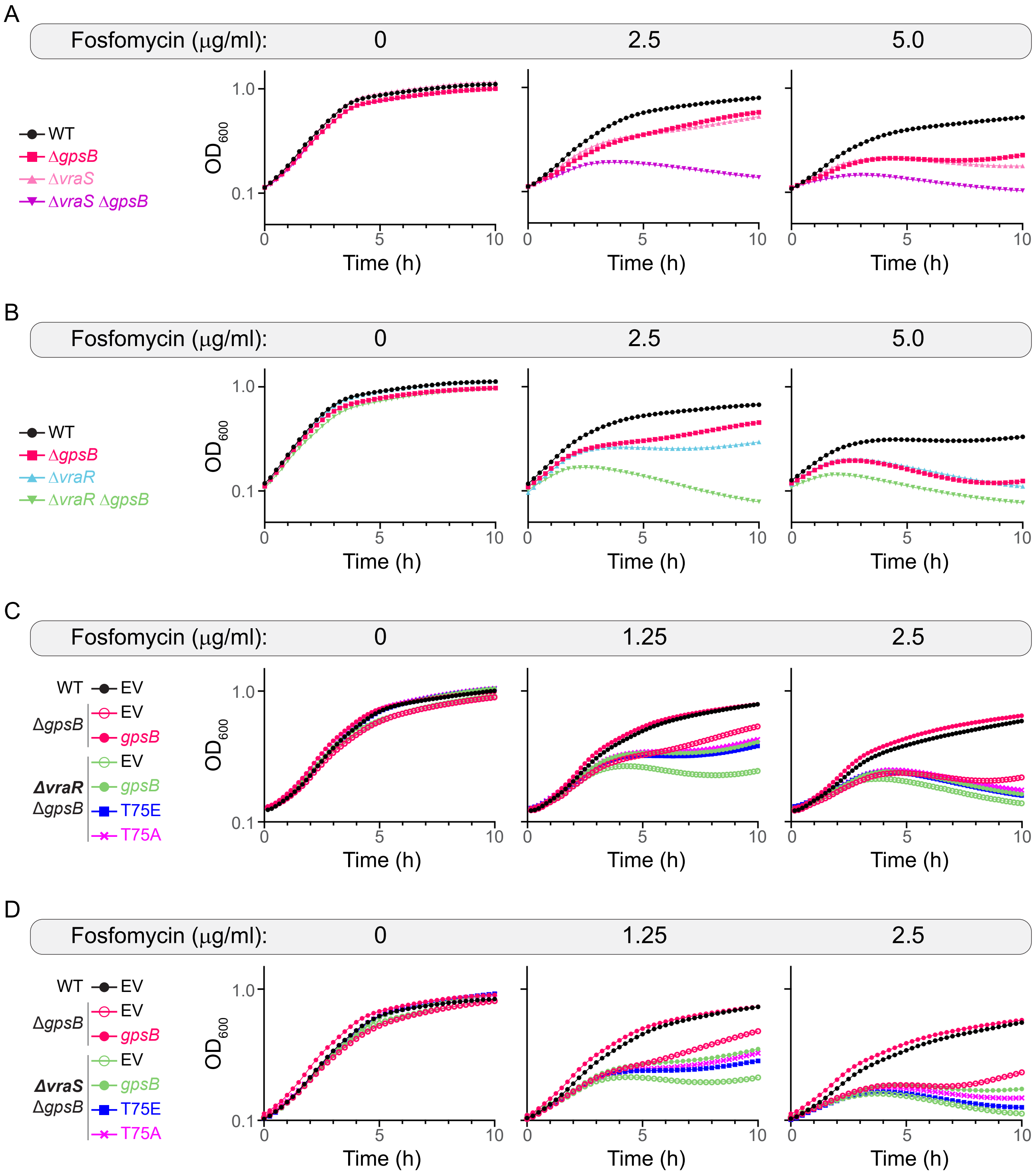
A
Fosfomycin (μg/ml): 0 2.5 5.0
WT
ΔgpsB
ΔvraS
ΔvraS ΔgpsB
OD600
Time (h)
B
Fosfomycin (μg/ml): 0 2.5 5.0
WT
ΔgpsB
ΔvraR
ΔvraR ΔgpsB
C
Fosfomycin (μg/ml): 0 1.25 2.5
WT EV
ΔgpsB EV
gpsB
ΔvraR ΔgpsB EV
gpsB
T75E
T75A
D
Fosfomycin (μg/ml): 0 1.25 2.5
WT EV
ΔgpsB EV
gpsB
ΔvraS ΔgpsB EV
gpsB
T75E
T75A

Figure S4
